# NMDAR-mediated transcriptional control of gene expression during the development of medial ganglionic eminence-derived interneurons

**DOI:** 10.1101/2020.06.10.144295

**Authors:** Vivek Mahadevan, Apratim Mitra, Yajun Zhang, Xiaoqing Yuan, Areg Peltekian, Ramesh Chittajallu, Caroline Esnault, Dragan Maric, Christopher Rhodes, Kenneth A. Pelkey, Ryan Dale, Timothy J. Petros, Chris J. McBain

## Abstract

Medial ganglionic eminence (MGE)-derived parvalbumin (PV)+, somatostatin (SST)+ and Neurogliaform (NGFC)-type cortical and hippocampal interneurons, have distinct molecular, anatomical and physiological properties. However, the molecular mechanisms regulating their diversity remain poorly understood. Here, via single-cell transcriptomics, we show that the obligate NMDA-type glutamate receptor (NMDAR) subunit gene *Grin1* mediates subtype-specific transcriptional regulation of gene expression in MGE-derived interneurons, leading to altered subtype abundances. Notably, MGE-specific conditional *Grin1* loss results in a systemic downregulation of diverse transcriptional, synaptogenic and membrane excitability regulatory programs. These widespread gene expression abnormalities mirror aberrations that are typically associated with neurodevelopmental disorders, particularly schizophrenia. Our study hence provides a road map for the systematic examination of NMDAR signaling in interneuron subtypes, revealing potential MGE-specific genetic targets that could instruct future therapies of psychiatric disorders.

## 1. INTRODUCTION

Medial ganglionic eminence (MGE)-derived forebrain GABAergic interneurons comprise the parvalbumin-containing (PV) and somatostatin-containing (SST) subpopulations throughout the entire forebrain accounting for approximately 60% of all cortical interneurons [1, 2]. In addition, approximately half of all hippocampal neurogliaform-type cells (NGFCs), the so called Ivy cells, originate from the MGE [3, 4]. Interestingly, though only rarely found in rodent neocortex such MGE-derived NGFCs are significantly more populous in primate neocortex, including humans [5]. While PV neurons exert robust somatic, and proximal dendritic inhibition, the SST and NGFCs mediate domain-specific dendritic inhibition on their downstream pyramidal neuron targets [6]. Collectively these classes of interneurons shape diverse aspects of cortical and hippocampal circuit maturation during development, and critically regulate information processing in mature circuits by maintaining appropriate excitation-inhibition (E-I) balance [7]. Recent evidence indicates a critical role for activity, particularly through ionotropic glutamate receptors (iGluRs), in driving the morpho-physiological maturation of MGE-derived interneurons [8–12]. Unlike mature interneurons where iGluRs differentially contribute towards synaptic transmission, immature and migrating interneurons express different glutamate receptor subunits including the NMDA-type iGluR (NMDAR) and AMPA/Kainate-type iGluR (AM-PAR/KAR) [13–15] prior to the expression of any functional synapses. This becomes particularly important as the developing brain contains higher ambient glutamate levels than the adult brain [16]. Collectively, higher ambient glutamate, developmental expression of iGluRs and recruitment of glutamatergic signaling is considered to be trophic [8, 17, 18] and thought to engage mechanisms to regulate various aspects of interneuron development including morphological and electrical maturation to promote appropriate circuit integration [9, 11, 14, 16, 19–22].

Interneuron-specific impairments are increasingly considered central to the etiology of multiple neurodevelopmental and circuit disorders [23]. The importance of interneuron-expressed iGluRs is most notable in psychiatric disorders exhibiting impaired NMDAR-associated systems [24, 25]. In the adult brain, acute pharmacological NMDAR blockade results in circuit disinhibition and psychotic symptoms [26], mediated in-part, by the enhanced sensitivity of interneuronal NMDARs to their antagonists [27]. Indeed, direct blockade of interneuron activity also precipitates distinct behavioral deficits relevant to schizophrenia [28]. In particular, ablation of the obligate NMDAR subunit gene *Grin1* in interneuron-specific early postnatal mouse [29], but not PV-specific [30], or glutamatergic neuron-specific *Grin1* ablation [31], resembles global *Grin1*-mutants [32] in their constellation of schizophrenia-like behavioral aberrations. This indicates that *Grin1* dysfunction across multiple interneuron-subtypes precipitates schizophrenia-like abnormalities [33]. In addition, this interneuron-specific NMDAR-hypofunction model is sensitive to developmental age, since adult-onset *Grin1* loss does not result in the same phenotypes [29]. Despite the importance of developmental NMDAR function in interneurons, and its relevance to human neurodevelopmental disorders, a comprehensive interrogation of the impact of developmental NMDAR ablation in interneurons, particularly across MGE-derived interneurons, is lacking.

It is clear that during the developmental window between embryonic day (ED) 13.5 and postnatal day (PD) ~10 [34], a combination of innate genetic programs, external environment, and neuronal activity shapes interneuron subtype specification leading to remarkable diversity [2, 21, 35, 36] The NMDAR signaling complex comprises an essential node for regulating gene expression via excitation-transcription (E-T) coupling in mature circuits [37–39]. Moreover, different NMDAR subunits are widely expressed in the developing brain [40] where they provide a critical source of Ca^2+^-entry via trophic glutamate signaling prior to synaptogenesis [15, 19, 41]. However, it is not clear whether the NMDAR-mediated Ca^2+^ cascades in nascent and developing MGE-derived interneurons engage transcriptional programs necessary for MGE-derived interneuron diversity. To investigate this, we conditionally deleted *Grin1* in MGE progenitors that give rise to cortical and hippocampal PV, SST, and NGFC subsets, using the *Nkx2-1*-Cre mouse line [3, 4, 42]. In this model, *Nkx2-1*-driven Cre expression is reported in cycling/proliferating MGE cells, well before the cells become postmitotic, allowing for assessment of the developmental impact of embryonic loss of *Grin1* activity across all subsets of MGE-derived interneurons. Applying a combination of high-throughput single-cell RNA sequencing (scRNAseq), MGE-interneuron-specific ribosomal tagging, and quantitative immunostaining and *in situ* RNAscope analyses we establish that NMDAR-mediated transcriptional cascades regulate MGE subtype abundances, by regulating the expression of diverse transcriptional, synaptogenic and membrane excitability genetic programs. Notably, we identify numerous diseaserelevant genes that are misexpressed in MGE-derived interneurons upon *Grin1*-ablation, providing a broad road map for examination of MGE-subtype specific regulation via NMDAR signaling.

## 2. RESULTS

### scRNAseq recapitulates cardinal MGE subtypes and a continuum of molecular profiles

To examine the molecular heterogeneity of MGE-derived GABAergic interneurons by scR-NAseq, we microdissected frontal cortex (CX) and hippocampus (HPC) from fresh brain slices obtained from PD18-20 *Nkx2.1*-Cre:Ai14 mouse **(Figure1A, Figure1-Supplement1A)**. Ai14-TdTomato (TdT^+^) single-cell suspensions were harvested by fluorescence-activated cell sorting (FACS) using stringent gating constraints including viability and doublet discrimination **(Figure1-Supplement1B)** as previously described [43–45], and subsequently processed through the 10X Genomics Chromium controller. 9064 and 9964 TdT^+^cells were recovered from cortex and hippocampus respectively across 3 biological replicates. To minimize the effect of excitotoxicity and stress-related transcriptional noise, the tissue processing, FACS, and sample collection steps were performed in buffers supplemented with Tetrodotoxin (TTX), DL-2-Amino-5-phosphonopentanoic acid (APV) and Actinomycin-D (Act-D) [46]. Because we observed concordant cell clustering across the replicates during preliminary analysis by Seurat v3 [47, 48] **(Figure1-Supplement2A)**, the replicates were pooled for in-depth analysis. Subsequent clustering and marker gene analyses revealed that ~ 62% and 33% of the TdT^+^ MGE-sorts from cortex and hippocampus respectively, express classical GABA markers including *Gadl / Gad2, Lhx6*; and the MGE-subclass markers *Pvalb, Sst,* and *Lamp5,* marking PV and SST, NGFC subsets respectively **(Figure1B, Figure1-Supplement3A)**. While we did not recover cells expressing the CGE-markers *Proxl*, *Htr3a* or *Vip,* we recovered a minor fraction of cells corresponding to glutamatergic neurons, astrocytes and microglia. In addition, ~25% and 71% TdT^+^ MGE-sorts from cortex and hippocampus respectively were enriched in oligo-dendrocytes marked by *Olig1* expression across all replicates **(Figure1-Supplement 2B,2C)**. However, we focused our subsequent analyses on the 5656 and 3002 *Gad1 / Gad2* positive cortical and hippocampal MGE-derived interneurons.

**Figure1.**
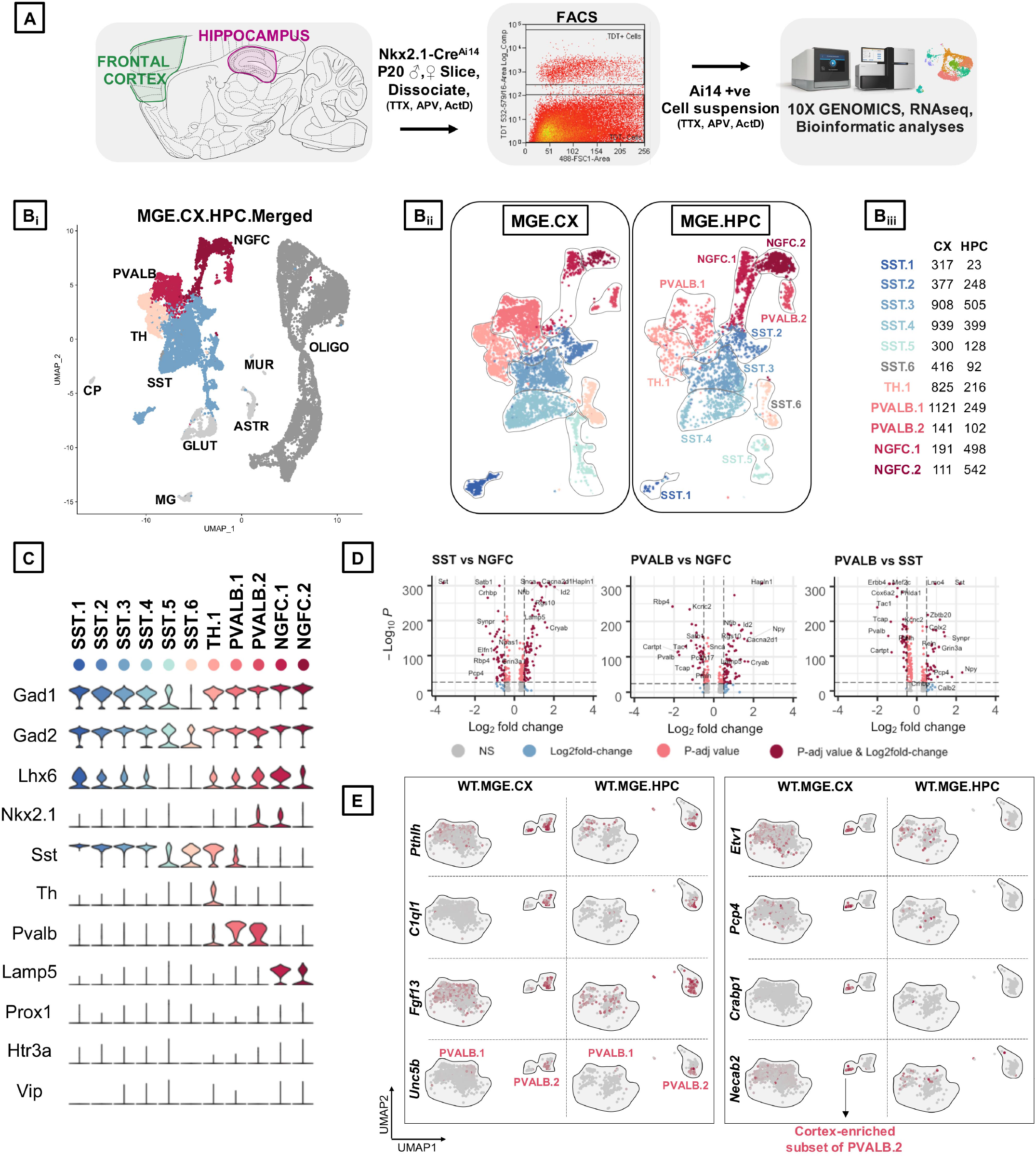
Identification of MGE-derived interneuron subtypes in the cortex and hippocampus. **A,** Overview of the experimental workflow. **B**¿, Uniform Manifold Approximation and Projection (UMAP) dimensional reduction of 19,028 single-cell transcriptomes (9,064 from frontal cortex and 9,964 from hippocampus of 6 mouse brains, 3 biological replicates), showing the cardinal MGE populations. Cell clusters were color coded and annotated *post hoc* based on their transcriptional profile identities (Cell type abbreviations: PVALB, Parvalbumin; NGFC, Neurogliaform; TH, Tyrosine Hydroxylase; SST, Somatostatin; GLUT, Glutamatergic; CP, Choroid Plexus; MG, Microglia; ASTR, Astrocyte; MUR, Mural; OLIGO, Oligodendrocyte). **B**_ii_, UMAP visualization of 11 MGE-derived interneuron subtypes from cortex (MGE.CX) and hippocampus (MGE.HPC), and the recovery of cell numbers from the subtypes. **B**_iii_, Table indicating the number of *Gad1/Gad2^+^cells* recovered in each MGE subtype from the cortex and hippocampus, and the defining genes enriched in each subtype. **C,** Violin plot showing the distribution of expression levels of well-known representative celltype-enriched marker genes across the11 MGE subtypes. **D,** —log10 False Discovery Rate (FDR) versus log2 fold change (FC) between each of the MGE cardinal class, representing the top enriched markers at a fold change ≥0.5 and FDR <10e-25. **E,** UMAP representation of PVALB clusters highlighting the cortex-specific enrichment of *Pthlh*-expressing PVALB.2 subtype that is not observed in the hippocampus.

Unbiased cell clustering by Seurat v3 identified six subtypes of SST, two subtypes of PV, two subtypes of NGFCs, and one subtype of Tyrosine hydroxylse (TH) expressing interneurons, expressing the markers Sst, *Pvalb, Lamp5* and *Th* respectively, across the two brain regions examined **(Figure1C)**. Notably, all but two subtypes (SST.5 and SST.6) expressed high levels of *Lhx6*, and 2 clusters corresponding to PV.2 and NGFC.1 expressed *Nkx2.1* at this developmental time. While the PV-SST- and NGFC-clusters clearly exhibited robust gene expression differences among each other **(Figure1D)**, the TH cluster appeared to express genes that correspond to both PV: SST clusters, including *Sst* and *Pvalb* expressions **(Figure1C, Figure1-Supplement3B)**. Particularly, at this developmental window we could not observe robustly different gene expression variances between the cortical and hippocampal counterparts, barring a few marginal, but significant differences **(Figure1-Supplement4B).** This gave us sufficient rationale to perform subsequent analyses using the MGE-derived interneurons pooled from cortex and hippocampus.

Among the **SST sub clusters**, SST.1-5 uniquely expresses *Chodl, Igf2bp3, Cdh7, Pld5* and *Nfix* respectively, while SST.6 expresses only markers that are common with other SST clusters **(Figure1-Supplement3B)**. With the exception of SST.6 the remaining SST-expressing subclusters are described in previous scRNAseq assays **(Figure1-Supplement5A)**. For example, the *Chodl*-expressing SST.1 cluster co-expresses high *Nos1, Tacr1, Penk,* and *Npy*, and it has been previously described as putative GABAergic long-range projection neurons [49, 50]. Clusters SST.2/3/4 express *Elfnl, Reln* and *Grm1* characteristic of putative cor-tical martinotti and their hippocampal counterpart, oriens-lacunosum/moleculare (O-LM) cells [43, 51, 52]**(Figure1-Supplement5B)**. Lastly, *Zbtb20*-expressing SST.5 is predicted to be septal-projecting interneurons [43]. Among the **PV sub clusters**, while both PVALB.1&2 coexpresses several common markers including *Pvalb*, *Kcnip2*, *Tcap* and *Kcnc1* there are several notable differences between the two clusters. PVALB.1 appears to contain continuous, but non-overlapping populations expressing *Syt2* representing putative fast-spiking basket cells or *Rbp4/Sst* containing putative bistratified cells [1, 43, 53]**(Figure1-Supplement5D)**. PVALB.2 contains cells that uniquely expresses *Pthlh, Clql1, Fgf13* and *Unc5b* representing putative axo-axonic chandelier cells [43, 49, 54]. We also observed a TH cluster, which, in addition to expressing several genes common to the SST: PV clusters, expresses several unique genes including *Rasgrp1*, *Bcl6*, *Myo1b* that segregated into mutually exclusive cluster space expressing *Crh* or *Nr4a2* **(Figure1-Supplement3B, Figure1-Supplement5B)**. This cluster is also described previously as putative bistratified-like cells [43, 53]. Among the **NGFC sub clusters**, while both NGFC.1&2 coexpress several common markers including *Lamp5, Hapln1, Cacna2d1, Sema3c* and *Id2,* the NGFC.1 cluster uniquely expresses several genes like *Reln, Ngf, Egfr, Gabra5* that are not expressed by NGFC.2. **(Figure1-Supplement3B)**. While the *Reln**+*** represents MGE-derived neurogliaforms, the Reln-population may represent putative ivy cells [43] **(Figure1-Supplement5C)**.

While the majority of the UMAP space aligns well between the cortical and hippocampal MGE-derived interneurons, we observed some regional differences as well. (**Figure1-Supplement3A**_i_**A**_iii_). (i) First, we observed an increase in the HPC-expressed NGFC.1&2 in comparison to their cortical counterparts, consistent with preferential localization of MGE-derived NGFCs to HPC over CX in rodents [1, 3, 4]. (ii) Next, the *Pthlh*-expressing PVALB.2 subcluster splits into two islands, only in the cortex and distinctly lacking from the hippocampus. Only one of the PVALB.2 islands expresses *Clql1*, while the other cortex-enriched island expresses unique markers *Etv1, Cnr1, Pcp4, Crabp1, Necab2, Epha4* and *Hapln1* **(Figure1E, Figure1-Supplement5D)** Whether this represents a novel subclass of chandelier cells remains to be determined. (iii) Lasty, we also observed a distinction in the hippocampal SST.3 corresponding to a subset of O-LM interneurons (**Figure1-Supplement3A_ii_**). The overall MGE cell numbers indicate that the SST cells account for the majority of MGE cell population recovered in the scRNAseq assay from both brain regions (Figure1-Supplement3A_ii_, **A_iii_**). The PV and TH clusters accounted for a greater share of MGE-derived interneurons in the CX than in the HPC. While it is plausible these relative cell proportions may be skewed by differential survivability of these subtypes during tissue dissociation, sorting and single-cell barcoding, these relative percentages were similar across biological replicates.

### NMDAR signaling maintains MGE-derived interneuron subtype abundance

Because neuronal activity and glutamatergic signaling are known to regulate multiple facets of interneuronal development [2, 21] [11, 36, 55], we hypothesized that the key obligate subunit *Grin1* and the NMDAR signaling complex may play an instructive role in determining MGE subtype identities. To test whether NMDAR signaling impacts the development and function of MGE-derived interneurons, we ablated them in MGE progenitors by crossing floxed-*Grin1* mice with the *Nkx2.1^Cre^* mouse line [42] **(Figure1-Supplement1A).** The earliest expressions of *Nkx2.1* and *Grin1* in the developing rodent brains is reported around ~embryonic day (ED) 10.5 and ~ED14 respectively [56–58]. Moreover, NMDAR-mediated Ca^2+^ signaling in migrating interneurons is reported by ~ED16 [15]. Because the expression and activity of *Nkx2.1* precedes *Grin1* expression, we rationalized that utilizing *Nkx2.1^Cre^* mouse will ablate *Grin1* and NMDAR signaling in MGE progenitors from the earliest developmental point. We sorted TdT^+^cells from the cortex and hippocampus of *Nkx2.1^Cre^*: *Grin1^f f^* :Ai14 mice and performed scRNAseq using the 10X platform. The scRNAseq experiments were performed using juvenile mice (PD18-20) of both sexes and from the same litters as the wildtypes (WT) to enable subsequent direct comparison. Similar to the WT-datasets, the MGE-*Grin1* mutants also revealed an enrichment of TdT^+^ oligodendrocytes **(Figure2-Supplement4B)**, however, we again focused our attention on the *Gadl/2* positive in-terneurons.

We next performed integrated analyses of the MGE-*Grin1^wt^* and MGE-*Grin1^fl/fl^* cortical and hippocampal scRNAseq datasets. Applying similar unbiased clustering parameters used for the MGE-*Grin1^wt^*analyses, we observed a total of twelve *Gad1/2* positive clusters in the integrated dataset **(Figure2A, B).** As a robust control, *Grin1* appeared to be absent or vastly reduced in all MGE subsets in both brain regions from MGE-*Grin1***(Figure2C),** but not in the *Slc17a7* expressing glutamatergic neurons **(Figure2-Supplement4A)**. Overlaying the WT and NULL datasets from the brain regions revealed differential enrichments among the recovered cells between the genotypes **(Figure2D)**. Intriguingly, *Grin1*-ablation did not seem to alter the SST or PV recovery percentages, with the exception of a modest increase in the cortical NGFCs (χ^2^ = 11.6, p = 0.003), but not hippocampal NGFCs (χ^2^ = 4.07, p = 0.13) (**Figure2E**_i_, **Figure2-Supplement2A_i_,B**).

**Figure2.**
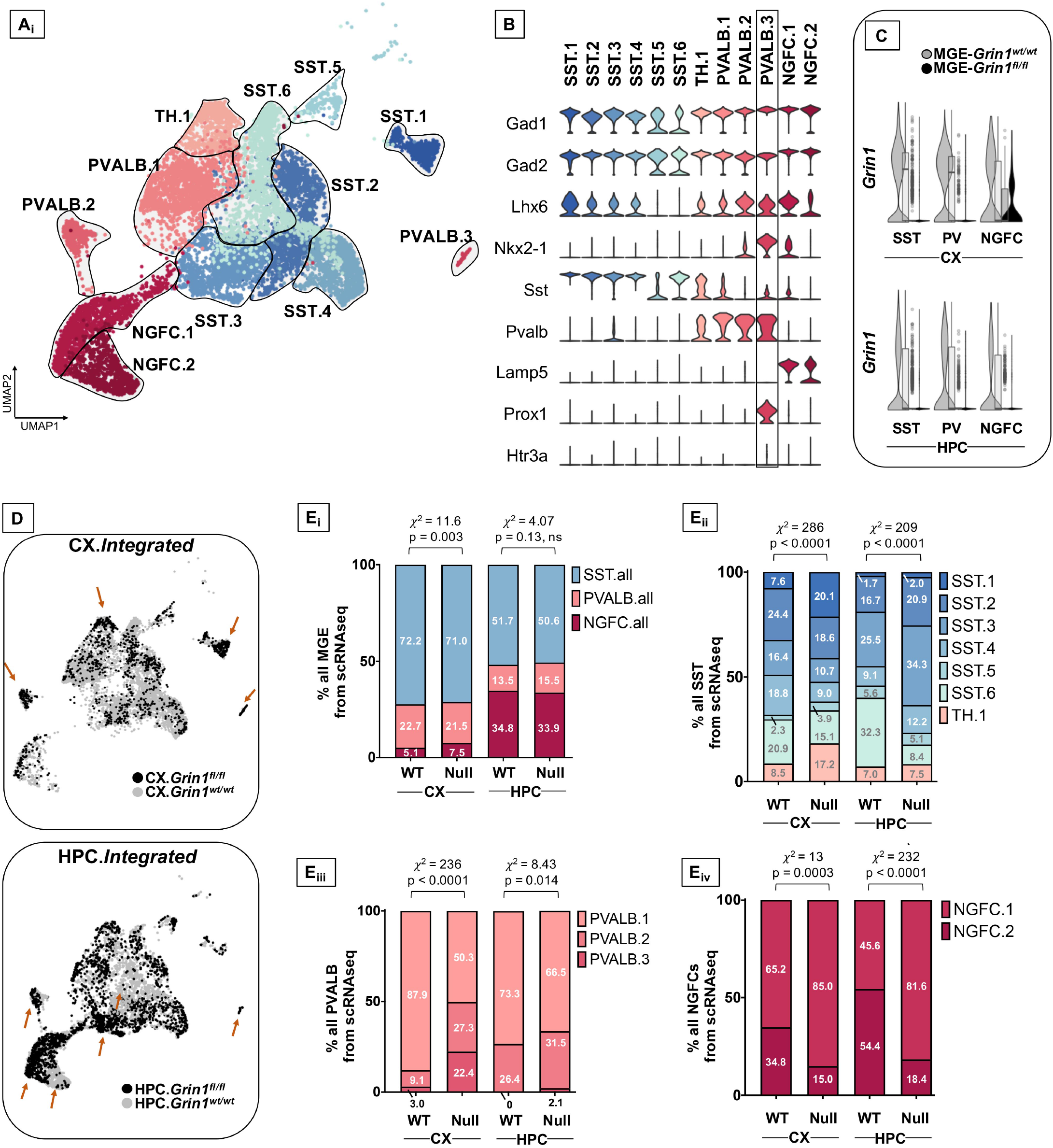
Altered interneuron subtype proportions upon *Grin1*-ablation. **A,** Integrated UMAP visualization of 12 subtypes of MGE-derived interneurons obtained from cortex (CX) and hippocampus (HPC) of *Grin1^wt/wt^* and *Grin1^fl/fl^* mice. UMAP represents the following numbers of MGE-derived interneurons from *Grin1^wt/wt^* and *Grin1*^fl/fl^respectively: 5624 (CX.WT) and 1387 (CX.NULL); 2998 (HPC.WT) and 2309 (HPC.NULL). 3 and 2 independent biological replicates of the scRNAseq assay from *Grin1^wt/wt^* and *Grin1^fl/fl^* respectively. **B,** Violin plot showing the distribution of expression levels of well-known representative cell-type-enriched marker genes across the 12 interneuron subtypes. **C,** Violin plot from both genotypes indicating the expression of *Grin1* in the cardinal of MGE-derived interneuron subtypes. **D,** UMAP representation colored by brain-region, highlighting the differential enrichments of cells (brown arrows) within interneuron subsets in *Grin1*-WT and *Grin1*-null from CX and HPC. **E,** Stacked-barplots representing the proportions of recovered cell numbers within **E**_*ii*_, pooled cardinal MGE subtypes, **E**_*ii*_, SST subtypes; **E**_*iii*_, PVALB subtypes and **E**_*iv*_, NGFC subtypes in *Grin1*-WT and *Grin1*-null from cortex or hippocampus. χ^2^, Chi-square test of proportions; ns, not significant.

To independently examine whether *Grin1* ablation impacts interneuron abundances, we conducted immunostaining experiments to probe the PV and SST subtypes from postnatal days (PD)30 brains from both genontypes. First we observed no change in the total TdT^+^ cell counts from both cortex and hippocampus (**Figure2-Supplement1 A**_*i*_, **B**_*i*_). Next, while we observed no change in hippocampal expressed total PV/SST cell type counts at PD30 (**Figure2-Supplement1B**_*ii*_), we observed a modest reduction in cortical PV cell type counts along with an increase in cortical SST cell type counts at the same age (**Figure2-Supplement1A**_*ii*_). This indicated differential impact of *Grin1*-ablation on cortical and hippocampal interneurons.

Despite observing no major changes in the recoveries of cardinal MGE-interneuron subtypes by scRNAseq, we observed marked changes in the recovery percentages of the subsets of SST, PV and NGFCs from both cortex and hippocampus (**Figure2E**_*ii*–*iv*_, **Supplement2A**_ii_**,C)**. Particularly, we observed a robust increase in cortical *Chodl*-expressing cortical SST.1 population, hippocampal *Reln*-expressing SST2-4 populations, and a decrease in hippocampal SST.6 population in MGE-Grin1^fl/fl^(CX, HPC: χ^2^ = 286, 209; p-value = 2.2e-16 for both regions). In addition, we found a reduction in the cortical PVALB.1 population, and a compensatory increase in PVALB.2/3 populations in MGE-*Grin1^fl/fl^* (CX, HPC: χ^2^ = 236, 8.4; p-value = 2.2e-16, 0.14). Finally, we observed an increase in the NGFC.1 along with a compensatory decrease in NGFC.2 in both cortex and hippocampus (CX, HPC: χ^2^ = 13, 232; p-value = 0.0003, 0.14). Among the differentially enriched subclusters, *Pthlh*-expressing PVALB.3 is quite notable **(Figure2B, Figure2-Supplement3A,B)**. This cortex-enriched cluster lacking in the hippocampus was identified within the PVALB.2 putative-chandelier cells in the MGE-*Grin1^wt/wt^* **(Figure1E, Figure1-Supplement5D)**, However, subsequent to integration of the MGE-*Grin1^fl/fl^* scRNAseq dataset, it segregated as a unique cluster, far from other PVALB clusters in the UMAP space. We observed *Proxl* expression in PVALB.3, which is uncharacteristic of MGE-derived interneurons, additional to robust expressions of genes associated with NGFCs such as *Hapln1* and *Reln* **(Figure2-Supplement3A,B)**. Moreover, we observed an increase in recovery of the cortical PVALB.3 cell numbers, including the emergence of these cells in the hippocampus subsequent to *Grin1*-ablation **(Figure2-Supplement2A,B).**

To independently examine whether the predicted differences in cell recovery percentages among the cardinal MGE subtypes are true, we conducted RNAscope in *situ* hybridization assays from PD20-25 cortex and hippocampus from both genotypes. We particularly focused on the subtypes of SST interneurons, namely the high-*Nos1*-expressing *Chodl*-SST.1 and the *Reln*-expressing SST.2-4 subtypes. First, similar to our scRNAseq prediction, we observed a significant increase in cortical *Sst*^+^ cells that co-express *Nos1* and a concomitant reduction in the *Sst*^+^ cells that lack *Nos1* after *Grin1*-ablation in MGE-derived interneurons **(Figure2-Supplement1C**_*i*_, **C**_*ii*_). Next, similar to our scRNAseq prediction, we observed an increase in hippocampal *Sst*^+^ cells that co-express *Reln* (**Figure2-Supplement1D**_*i*_, **D_*ii*_**), without changes in total Sst+ cell numbers in both brain regions. It is unclear whether the changes in subtype numbers reflect a change in cell identity or whether this is alterered subtype proportions due to differential survival. Nevertheless, these data demonstrate clear changes in MGE-derived interneuron subtype abundances following early embryonic loss of *Grin1* function.

### NMDAR signaling shapes the transcriptional landscape in MGE-derived interneurons

It is now well-established that transcriptional signatures defines the subtype identities of GABAergic interneurons [59]. To examine the full range of transcriptional impairments triggered by *Grin1* ablation in MGE-derived interneurons, we next performed differential gene expression testing by pooling the SST / PVALB / NGFC subtypes into their cardinal MGE classes to identify the genes that are differentially expressed between the genotypes. For instance SST1-6 and TH.1 are pooled together as SST; PVALB1-3 are pooled together as PVALB, and NGFC1-2 are pooled together as NGFC cardinal classes for this assay. At a stringent false-discovery rate (FDR) <0.01, 802 genes passed the 10%-foldchange (FC) threshold across the MGE subtypes from both brain regions **(Figure3A, Supplemental Table1**). Several interesting features were observed in the differentially expressed gene (DEG) pattern upon MGE-specific *Grin1*-ablation. (i) Among all DEGs only ~10% and 1% are upregulated in the cortex and hippocampus respectively, while the remaining genes were all down-regulated (**Figure3-Supplement1A_*ii*_,B,C**). (ii) While *Grin1* ablation resulted in several unique DEGs between the MGE classes, ~10 and 27% of the DEGs are common within cortex and hippocampus respectively **(Figure3A, Figure3-Supplement2)**. For instance, while *S100a10, Hapln1, Hcrt2* are uniquely upregulated in cortical SST, PV and NGFC respectively **(Figure3A)**, *Apoe, Kcns3, Wnt5a* were uniquely altered in hippocampal SST, PV and NGFC respectively. In contrast, *Grin1* ablation induced common changes in *Penk1* and *Erbb4* expression patterns across all MGE-derived interneuron classes in the cortex and hippocampus respectively. (iii) ~27-43% of all DEGs were shared by MGE classes across brain regions **(Figure3-Supplement2A_*ii*_)**. For example, *Npas3, Cdh9, Grm1* are commonly downregulated in all SST subclasses; *Bcl6, Epha7, Gabra4* common to PV class; and *Rgs12, Gabrad, Sema5a* common to NGFCs from both brain regions. (iv) Lastly, 28 genes are commonly differentially expressed across both brain regions, across all MGE subtypes. For example, *Grin1, Neto1, Cdh2, Scn2b* are commonly downregulated across the board, while *Epha5, Olfm1* are commonly downregulated across all, but cortical PV cells **(Figure3A)**.

**Figure3.**
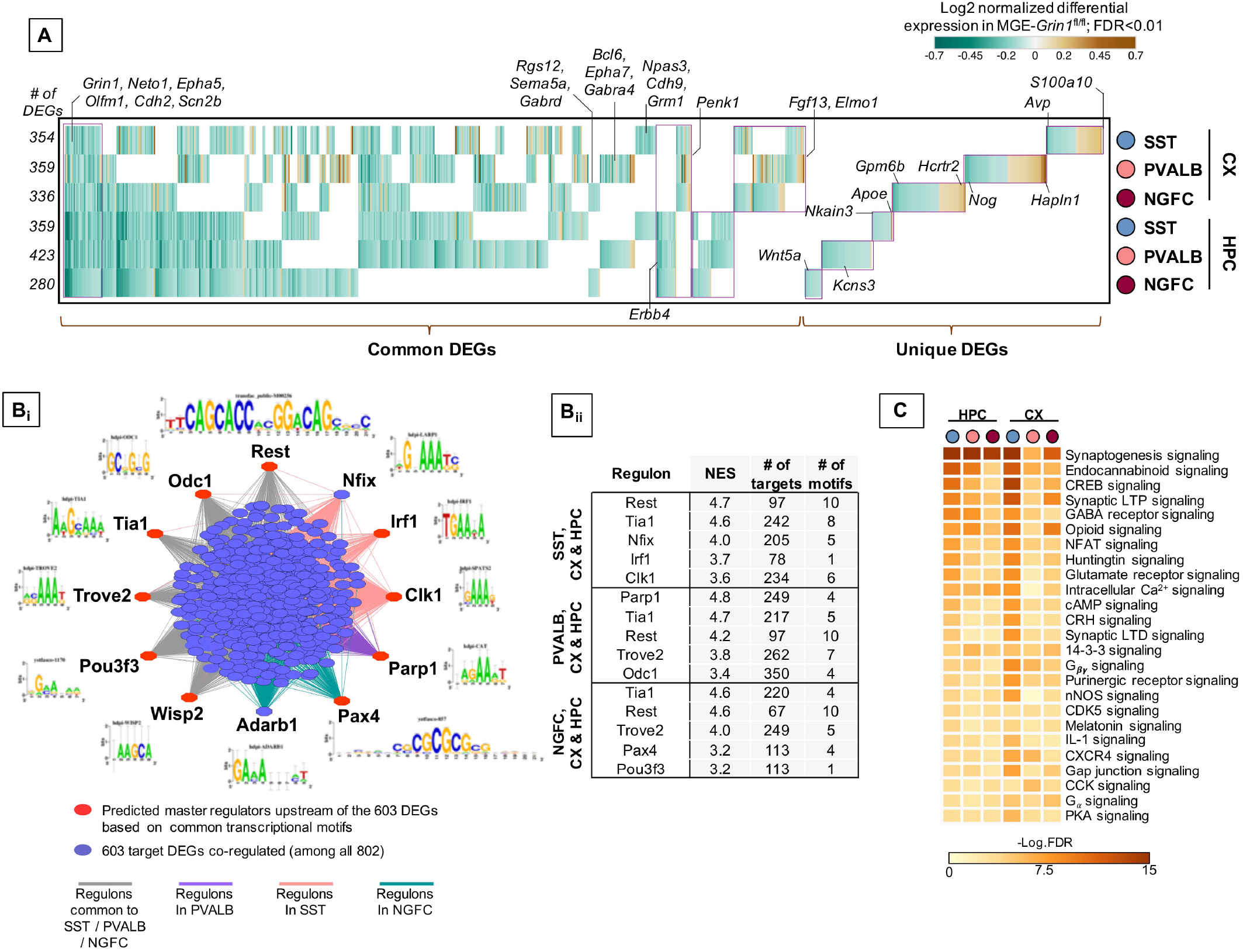
Cell-autonomous transcriptional changes subsequent to MGE-specific developmental *Grin1*-ablation. **A,** Combined heatmap representing the 802 differentially expressed (DEGs) in the cortical and hippocampal MGE-derived interneurons upon *Grin1*-ablation, at a FDR<0.01 and FC>10%, as determined by MAST analysis (see details in Methods). **B**_i_, iRegulon in *silico* analysis identifying high-confidence master upstream transcriptional regulators(indicated in red) of the DEGs (indicated in lavender). Representative DNA-binding motifs are indicated next to the transcriptional regulators. **B**_ii_, Top five transcriptional regulators predicted by iRegulon, associated normalized enrichment score (NES) and number of predicted targets and motifs associated with each transcription factor cluster, indicated for the three interneuron subtypes (CX and HPC pooled). **C,** Ingenuity Pathway Analysis of significantly overrepresented molecular pathways in each MGE-subtype. FDR determined by Fisher’s Exact Test.

Gene expression co-regulation is intrinsic to cellular diversity [60, 61]. Since the majority of DEGs are downregulated across the MGE subtypes, we examined whether they correspond to clusters of coordinated co-regulation. We applied the iRegulon *in silico* framework [62], which identifies transcription factor binding motifs that are enriched in genomic regions of the DEGs upon *Grin1*-ablation, and predicts the transcription factors that bind to the motifs. This *in silico* analysis predicted 51 significantly enriched motifs (normalized enrichment score > 3) that clustered into 10 groups by similarity, 33 of which were associated with transcription factors **(Figure3B**, **Supplemental Table2)**. Put together, 10 transcription factors were predicted to bind with the motifs with high confidence, strongly supporting targeted co-regulation of 617 among the 802 DEG genes upon *Grin1*-ablation. Notably, the RE1-silencing transcription factor (*Rest*) is a master transcriptional repressor that mediates the transcriptional accessibility for several synaptic genes [63], including NMDAR subunits themselves [64]. It is intriguing to observe that the downregulation of the DEGs upon MGE-specific *Grin1*-ablation are, in part, predicted to occur via *Rest*-mediated transcriptional repression.

To examine the broad biological impact of the DEGs, we performed Gene Ontology (GO) analyses. Broad GO analyses on all DEGs indicates that these genes serve to regulate multiple molecular functions in interneurons, including regulation of GABAergic and glutamatergic synapses, additional to biological pathways related to ad-diction and circadian entrainment **(Figure3-Supplement2B).** Further classification of DEGs based on their cellular functions within the MGE subtypes revealed genes critical for regulation of membrane excitability, gene expression, synaptic partnering and assembly, as well as major intracellular Ca^2+^ signaling cascades and second mes-sengers **(Figure3C, Figure3-Supplement2C, Supplemental Table3).**

### Transcription factor expression is a key component of NMDAR-mediated regulation of MGE-derived interneurons

Because transcriptional regulation underlies numerous fundamental processes including the expression of other classes of genes, we next examined the DE-transcriptional regulators in detail. We first examined the 67 genes that are differentially expressed upon *Grin1*-ablation and are known to mediate transcriptional regulation of gene expression. Of these, 35 genes are previously established to be expressed in different GABAergic interneuron classes including some notable MGE-expressed transcription factors **(Figure4A_*i*_)**. The remaining 32 are broadly ex-pressed TFs **(Figure4A_*ii*_)**, that include a small subset of 15 genes that are regulated by neuronal activity. Barring a few genes, we observed the majority of TFs to be down regulated in both brain regions. Intracellular Ca^2+^ signaling cascades and second messenger systems are key mediators of NMDAR signaling to the nucleus for transcriptional regulation. Theoretically, an early first wave impairment of Ca^2+^ signaling in *Grin1*-lacking MGE progenitors could result in transcriptional silencing of the mediators of Ca^2+^ signaling cascades and second messenger systems, which would sustain the transcriptional impairments. Indeed, we also observed a downregulation of various Ca^2+^ homeostasis-regulators, kinases / phosphatases and second messengers that are activated downstream of *Grin1* **(Figure4-Supplement1A,B,C)**. Furthermore, we noted that hippocampal MGE neurons had a greater proportion of DE-TFs and kinase signaling cascade effectors that were downregulated across all 3 subtypes compared to their cortical counterparts. Together, this suggests that hippocampal MGE-derived interneurons may be more vulnerable than cortical MGE-derived interneurons towards *Grin1*-mediated Ca^2+^ transcriptional silencing at this age.

**Figure 4.**
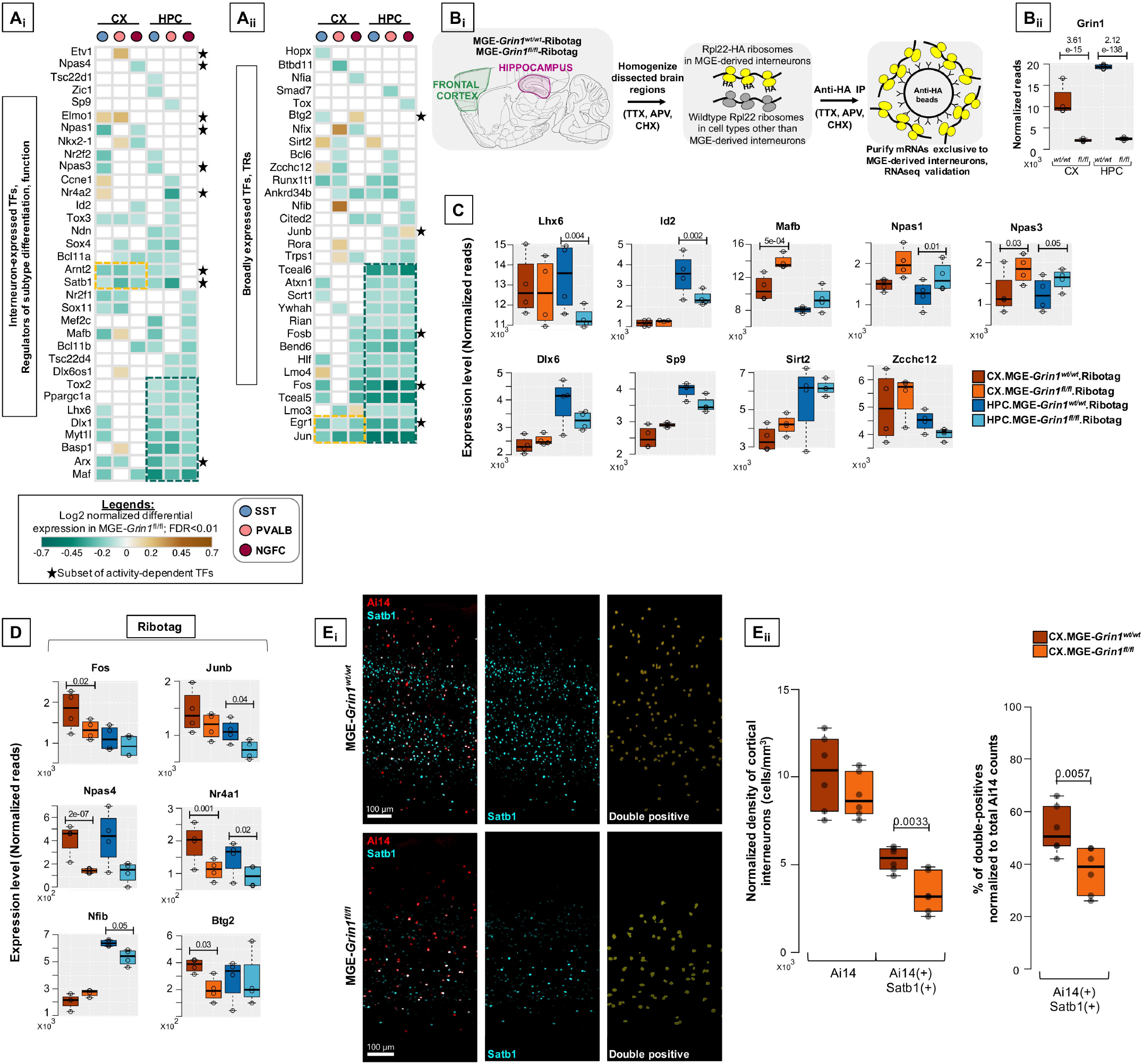
Grin1-signaling in MGE-derived interneurons are highly dedicated to the transcriptional control of interneuron identity. **A,**Heatmap of log2 FC of significant DEGs in cortical and hippocampal MGE cardinal subtypes, showing a subset of **(A**_i_) Transcription factors (TFs) that are previously established to regulate MGE subtype identity and function; **(A**_ii_) broadly expressed TFs and transcriptional regulators (TRs) that are not currently known to regulate MGE function. Clusters of commonly differentially expressed genes in cortex and hippocampus are indicated in yellow or green boxes. **\*** indicates a subset of neuronal activity-regulated TFs. **(B**_i_) Overview of the experimental workflow for RNAseq validation, by ribosomal tagging / Ribotag strategy specific to MGE-derived interneurons. n = 4 biological replicates from each genotype for MGE-Ribotag-seq; Error bars reflect s.e.m; DeSeq2 for statistical analysis. **(B**_ii_) MGE-Ribotag-seq normalized reads for *Grin1* expression in cortex and hippocampus. MGE-Ribotag-seq normalized reads cross-validating the differential expressions of **(C)** MGE-enriched TFs, and **(D)** neuronal activity-regulated TFs. **(E**_i_) Representative immunostaining of P25-30 somatosensory cortex using anti-Satb1 (cyan), and endogenous Ai14 reporter expression. Ai14:Satb1 doublepositive cells are represented as yellow surfaces using Imaris9.6. **(E**_ii_) Boxplots indicate the normalized density of cortical Ai14^(+)^ and Ai14^(+)^ :Satb1^(+)^ double-positive cell counts. n = 3 brains from each genotype for immunostaining; Error bars reflect s.e.m from individual sections; two-tailed unpaired *t*-test, for statistical analysis.

Interestingly, among the early TF cascades in the progenitors that sequentially determine and maintain MGE fate, several members appear to be expressed at ~P20, and starkly downregulated upon *Grin1*-ablation. For instance, *Lhx6, Maf, Arx, Myt1l, Dlx1* are among the genes broadly downregulated across all hippocampal MGE subtypes and within specific class(es) in their cortical parallels **(Figure4A)**. Other MGE fatedetermining TFs, *Nkx2-1, Mafb, Satb1, Nr2f1* (CoupTf1), *Sp9,* also appear to be downregulated in discrete populations. This also includes a downregulation of *Bcl11b* (Ctip2) in both hip-pocampal and cortical NGFCs, a gene recently linked to regulation of NGFC morphology and function [65]. Among the few transcriptional regulators upregulated are *Sirt2, Elmo1, Zcchc12,* none of which have been characterized in the context of MGE function **(Figure4B)**. *Sirt2* is an established transcriptional repressor [66, 67] that may regulate the repression of several target genes in an MGE-specific manner, and *Elmo1* has been previously characterized during the activitydependent migration of CGE subtypes [21]. Finally, a recent study has predicted that the expression of *Zcchc12* correlates with slower intrinsic firing among hippocampal CA1 interneurons [43]. This suggests that increased *Zcchc12* expression might regulate the expression of synaptic genes enabling reduced intrinsic excitability in the MGE subsets. Related to such putative decreased excitability in the MGE-derived interneurons, among the activity-regulated TFs, we observe broad downregulation of *Jun, Egr1, Fos, Fosb, Arc, Satb1, Arnt2* across all classes of MGE in both brain regions (**Figure4A,B**, indicated by *). While most of these are well-established activity-regulated TFs, *Arnt2* has been recently described to partner with *Npas4,* downstream of Ca^2+^ signaling in response to neuronal activity [68]. Unsurprisingly, *Npas4* is also significantly downregulated in discrete MGE subtypes.

### MGE-Ribotag-seq cross-validates impaired transcription factor expressions due to MGE-*Grin1*-ablation

To independently validate the changes in gene expressions observed by scRNAseq, we employed a Ribotag-based strategy where we generated a triple-transgenic mice by breeding the *Nkx2-1^cre^:Grin1^fl/fl^* with the *Rpl22^HA/HA^-* containing Ribotag mice [69], hereafter called MGE-Ribotag **(Figure4B**_*i*_**, Figure4-Supplement1D)**. We recently established that this is a robust tool to examine the mRNAs associated with the translational machinery, in a manner exclusive to MGE-derived GABAergic interneurons [70]. mRNAs copurified with anti-HA immunoprecipitation were utilized to prepare cDNA libraries that were subsequently sequenced in order to investigate changes in gene expression changes resulting from MGE-specific *Grin1*-ablation. By utilizing cortical and hippocampal tissues from age-matched MGE-Ribotag from both genotypes, we first established a robust reduction in *Grin1* expression in both brain regions (**Figure4B**_*ii*_). Similar to the observed changes in the scRNAseq assay we observed robust changes in several key TFs by MGE-Ribotag-seq, particularly in genes that have established functions in maintaining MGE-identities. Notably, we observed a significant downregulation of hippocampus-expressed *Lhx6* and *Id2*, an upregulation in cortical *Mafb* and hippocampal *Npas1*, and an increased expression of *Npas3* in both brain regions, after *Grin1* ablation **(Figure4C, top)**.

Key differences exist between the two strategies scRNAseq and MGE-Ribotag-seq. The scRNAseq assays only the mRNAs present in the cytoplasm of the neurons while providing a single-cell resolution across the subtypes of MGE-derived interneurons. The MGE-Ribotag-seq however lacks the single-cell resolution across MGE-interneuron subtypes but includes the mRNAs that are bound to the translational machinery and much closer to the cellular proteome. Moreover, MGE-Ribotag-seq includes the translating-mRNAs present in the neuronal processes. Despite these technical differences, the congruity in the DEGs after *Grin1*-ablation strongly favors our claims related to NMDAR-signaling dependent regulation of key transcription factors. Having a gene not validated by MGE-Ribotag-seq does not imply a false-positive scRNAseq result, and the above-described technical differences could contribute to it. Also, the genes that are commonly differentially expressed across the PV-SST-NGFC subtypes by scRNAseq will have a greater possibility of being identified as a DEG via MGE-Ribotag-seq. Accordingly, genes such as *Dlx6*, *Sp9*, *Sirt2* and *Zcchc12* that showed alternating differences among subtypes in the scRNAseq, did not reveal significant differences via MGE-Ribotag-seq **(Figure4C, bottom)**.

We also observed a significant decrease in the expressions of several activity-regulated TFs by MGE-Ribotag-seq in both brain regions **(Figure4D)**. While *Fos, Junb, Btg2, Nfib* expressions were significantly downregulated in either cortex or hippocampal MGE-derived interneurons, the expressions of *Nr4a1* and *Npas4* appear to be commonly downregulated in both brain regions after *Grin1*-ablation. *Satb1* is a key activity-regulated TF that is expressed in MGE-derived interneurons, critically regulating the identity and survival of different subtypes [71, 72]. Independent immunostaining experiments revealed that *Grin1*-ablation resulted in a robust reduction in MGE-derived interneurons co-expressing *Satb1* in PD30 somatosensory cortex (**Figure4E**_*i*_**, E**_*ii*_), further substantiating our findings from scRNAseq. Taken together, this establishes a framework for simultaneous regulation of neuronal activity and expression of distinct sets of TFs (including the activity-regulated TFs) by NMDAR-signaling in MGE-derived interneurons.

### Impaired NMDAR signaling alters region-specific MGE subtype marker expression

Several GABAergic/MGE markers were misregulated upon *Grin1*-ablation **(Figure5A)**. For example, genes *S100a0, Pthlh, Hcrtr2* that are normally expressed in SST, PV and NGFCs respectively, are upregulated in the same clusters of MGE-*Grin1*^fl/fl^**(Figure5B_*i*_**), indicating a misexpression in a subtype-specific manner. Next, while certain genes such as *Reln, Tenm1* are broadly downregulated across MGE classes, some genes like *Thsd7a* show an upregulation in certain classes but a downregulation in the other classes (**Figure5B**_*ii*_). Interestingly, a few genes that are normally abundant in one MGE class, appear to be misexpressed in another MGE class where they are not abundant. For instance, *Tcap*, that is normally expressed in PV cells, in addition to being decreased in PV cells, is upregulated in NGFCs in both cortex and hippocampus. Similarly, *Hapln1* expression which is typically limited to NGFCs, is upregulated in PV subsets (**Figure5B_*iii*_**). Indeed, MGE-Ribotag-seq cross-validates a robust downregulation in *Tcap*, *Tenm1* and *Hapln1*, and a robust upregulation in *S100a0, Tacr1* and *Nos1* **(Figure5C)**. Lastly, we observed an upregulation in the *Gad1* and *Slc32a1* (vesicular GABA transporter, vGAT) and a downregulation in *Gad2* and *Slc6a1* (Na^+^-Cl dependent GABA transporter, GAT1), corresponding with GABA synthesis and reuptake machineries respectively **(Figure5-Supplement5A)**. These findings closely match the trending differences observed via MGE-Ribotag-seq.

**Figure5.**
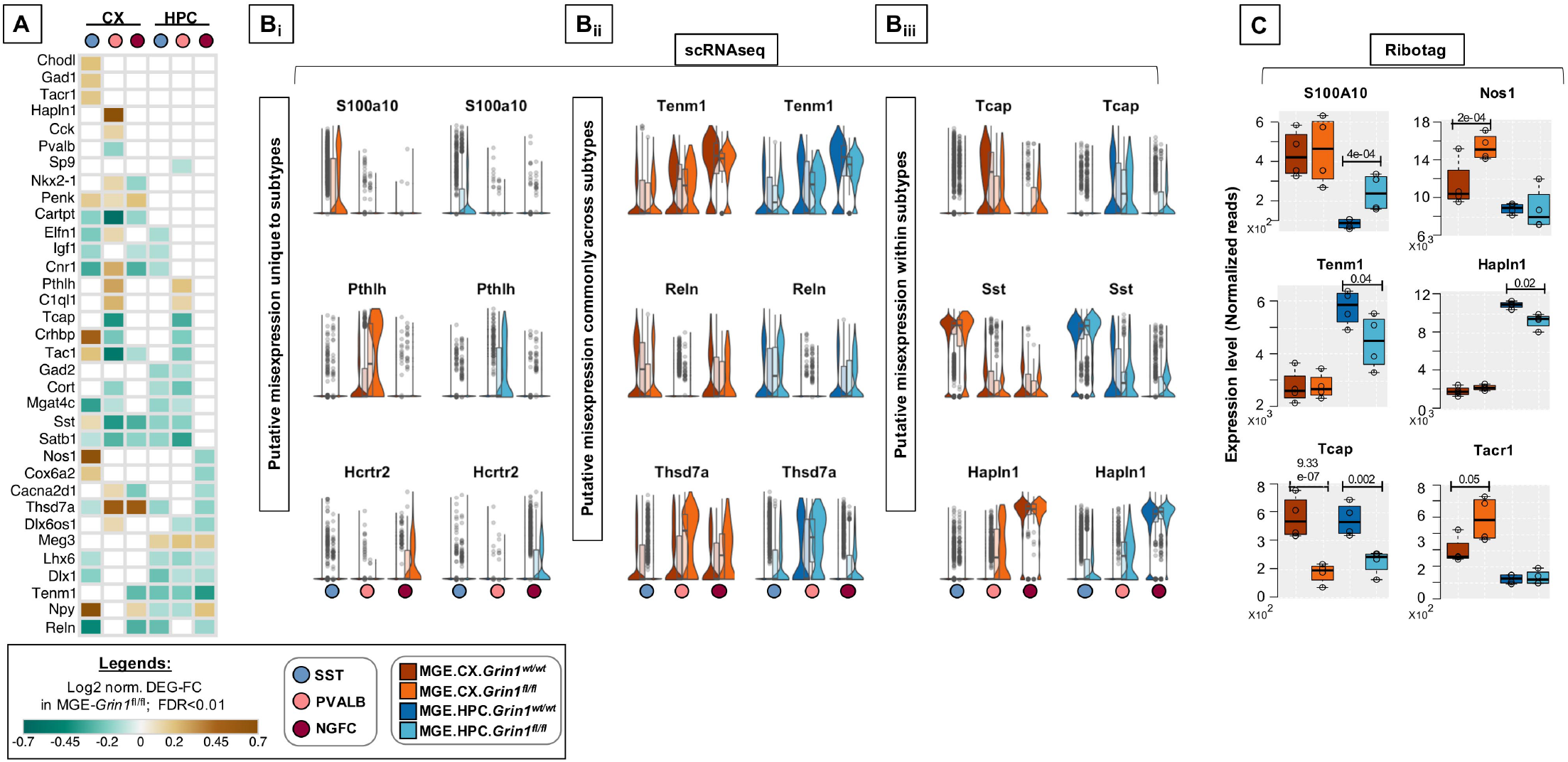
Differential expression of interneuron marker genes across subtypes upon *Grin1*-ablation. **A,** Heatmap of log2 FC of significant DEGs in cortical and hippocampal MGE cardinal subtypes, showing a subset notable MGE marker genes. Representative box-violin plots of top differentially expressed genes from the above that represent **B**_i_, a misexpression unique to a single MGE-*Grin1^fl/fl^* subtype; **B**_ii_, a misexpression across all MGE-*Grin1^fl/fl^* subtypes; **B**_iii_, a misexpression between MGE-*Grin1^fl/fl^* subtypes. **(C)** MGE-Ribotag-seq normalized reads, cross-validating the misexpressed interneuron marker genes in cortex and hippocampus.

### NMDAR signaling regulates MGE subtype-specific expression of neurodevelopmental disorder risk genes

Interneuron-centric disease etiology is an emerging centrality in multiple psychiatric disorders [23]. Thus, we questioned whether the *Grin1* ablation induced DEGs presently identified correlate with disease etiology. Disease-ontology based Fisher’s Exact Test conducted on the DEGs showed significant over-representation of genes implicated in ‘Schizophrenia’, ‘Psychiatric disorders’ and ‘Movement disorders’, among other cellular impairments involving aberrant morphology of neurons **(Figure6A, Supplemental Table4)**. To independently examine the DEGs for potential enrichment for neurodevelopmental disorders, we obtained the risk genes for schizophrenia (Sz) and autism spectrum (As) from the SZDB [73] and SFARI [74] databases respectively. These databases curate and rank disease-relevant gene sets, based on multiple evidence sources including genome-wide association studies, copy-number variations, known human mutations and other integrative analyses. In particular, we mapped the DEGs with the topranked genes from these disease datasets (see methods for details). While 592 DEGs could not be mapped with either disease genes, 25 genes mapped exclusively with the SFARI-AS gene list, 164 genes mapped exclusively with the SZDB-Sz gene list and 21 genes mapped with both datasets **(Figure6B_i_, Supplemental Table5)**. It is now well-established that several neurodevelopmental disorders exhibit a high degree of converging molecular pathways employing proteins that exist in physical complexes [75–78]. Therefore, we examined whether these disease-associated DEGs are known to form protein complexes between each other, by mapping curated protein-protein interaction (PPI) datasets for all 802 DEG products. Indeed, we observed that >95% of disease-annotated DEG products are known to exist with PPIs, while only ~75% of DEG products not annotated with Sz/As are known to exist with PPIs **(Supplemental Table5C)**. Interestingly, despite not mapping directly with the high-ranked disease gene sets, the remaining 592 genes are observed to exist in tightly-knit PPIs with the disease annotated genes. However, the PPIs mapped with SZDB form the most interconnected clusters in comparison to the SFARI-mapped PPI network (**Figure6B**_*ii*_), as indicated by relatively higher clustering coefficient. Put together, both unbiased disease ontology prediction, targeted analysis of high-confidence neuropsychiatric disorder genes, indicate that members of the DEGs share physical, and functional pathways in MGE-derived interneurons, contributing towards disease etiology.

**Figure6.**
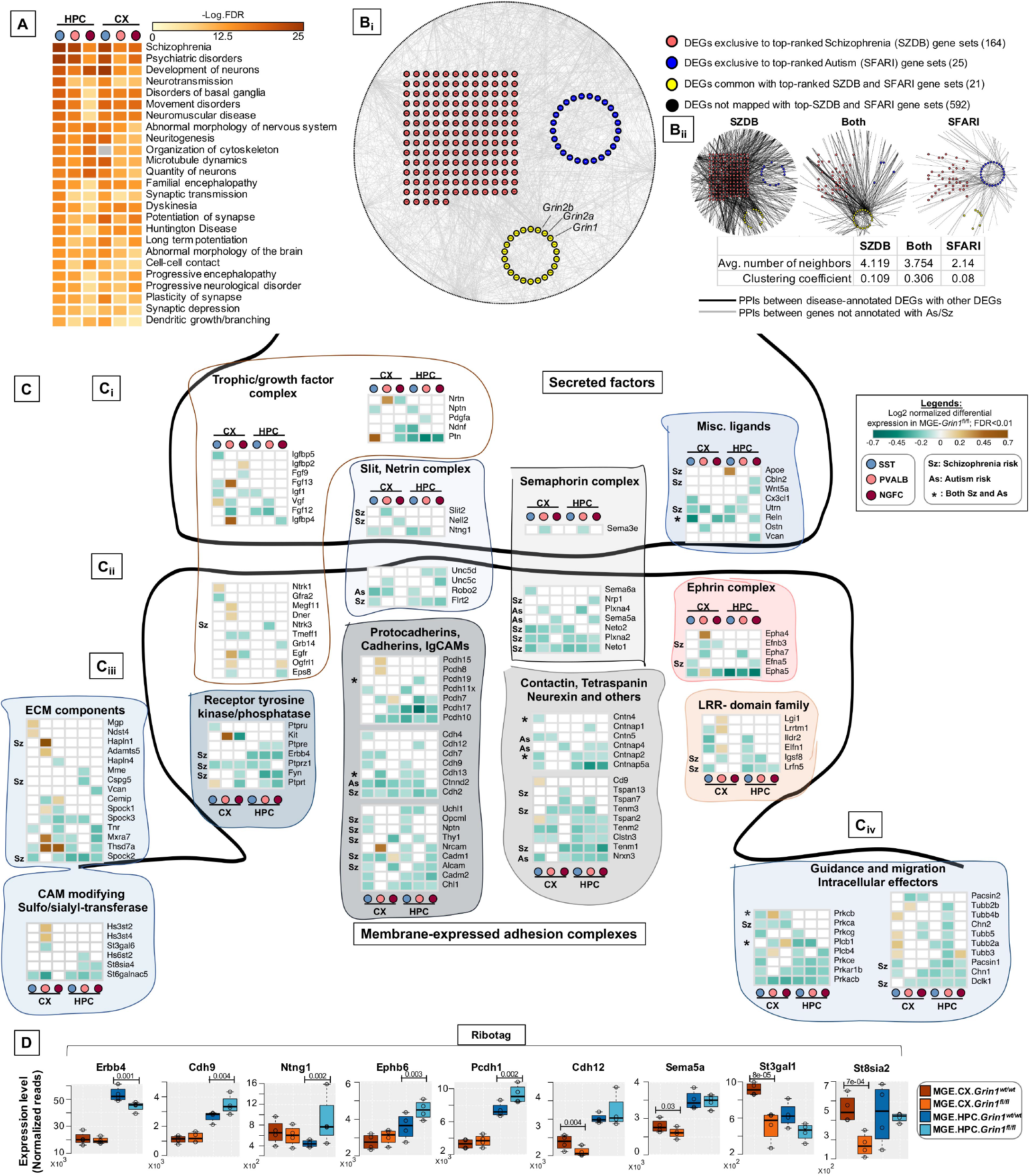
Aberrant *Grin1*-signaling result in misexpression of high-risk Sz genes. **A,** Ingenuity Pathway Analysis of significantly overrepresented disease pathways in each MGE-subtype. FDR determined by Fisher’s Exact Test. **B**_i_, Global protein-protein interaction (PPI)map among all differentially expressed genes (DEGs). Red circles indicate the DEGs annotated to be top-ranked Sz-risk genes; Blue circles indicate the DEGs annotated to be top-ranked As-risk genes; Yellow circles indicate the DEGs annotated with both Sz and As-risk genes. Black circles in the periphery indicate the DEGs not annotated with high-risk Sz/As genes. The PPIs between DEGs indicated in grey lines. **B**_ii_, PPIs between Sz / As / dually enriched clusters, and other genes. The PPIs between disease-annotated DEGs and other disease-annotated DEGs or with other non-annotated DEGs are indicated in black lines. ThePPI between nonannotated DEGs indicated in grey lines. **C,** Heatmap of log2 FC of significant DEGs in cortical and hippocampal MGE cardinal subtypes, showing a subset of **C**_i_, secreted trophic factors and secreted ligands and guidance cues. **C**_ii_, membranebound synaptogenic receptors and cell adhesion molecules (CAMs) **C**_iii_, Extracellular Matrix (ECM) components and matrix modifying enzymes. **C**_iv_, Intracellular effectors of guidance and synaptogenic cues. **D,** MGE-Ribotag-seq normalized reads, cross-validating the differentially expressed synaptic partnering molecules.

Among the 210 DEGs mapped with to Sz and As, 45 genes are established regulators of axon path-finding, synapse formation and assembly, while 38 members are established regulators of membrane excitability and neuronal firing. Because both of these gene classes are intimately associated with interneuron function, we examined these classes in detail. We observed multiple classes of secreted ligands and cognate receptor families corresponding to semaphorin, netrin, slit, chemokine and growth factors, and their intracellular effectors that are downregulated upon MGE-*Grin1*-ablation (**Figure6C**_i,iv_). These include *Ntng1, Sema3e, Slit2, Cx3cl1,* and some of their receptors, *Unc5c, Nrp1, Neto1/2, Robo2* that are decreased in a MGE-class-specific manner. We observed *Fgf13* that was recently demonstrated to mediate MGE-subtype specific synapse assembly [79], to be upregulated in cortical PV cells, but downregulated in cortical SST, while *Apoe* to be upregulated in hippocampal SST cells. In addition to synaptic assembly molecules, we observed DE in a variety of synaptic adhesion molecules, corresponding to protocadherin, cadherin, ephrin and contactin families (**Figure6C_ii_**). Notably, we also observed a downregulation of *Erbb4* across all hippocampal MGE-subtypes. Lastly, we observed increased expression of extracellular matrix components *Mgp, Ndst4, Hapln1, Adamts5, Mxra7, Thsd7a* and the matrix modifying enzymes *Hs3st2/4* in cortical SST/PV subtypes (**Figure6C**_iii_). In parallel, MGE-Ribotag-seq cross-validates a robust down regulation in hippocampal *Erbb4,* and a robust differential expression of several synaptic partnering and adhesion molecules including *Cdh9, Ntng1, Ephb6, Pcdh1, Cdh12, Sema5a*, and key members of CAM modifiers *St3gal1* and *St8sia2* **(Figure6D)**.

Among the regulators of neuronal excitability, we observed a downregulation of multiple members of postsynaptic glutamate receptor subunits, GABA receptors and their associated partners (Figure6-Supplement1B_ii_). Interestingly, while we noted a broad downregulation of several members of potassium and sodium channel sub-units, a few discrete members of the *Kcn*-families were upregulated in cortical PV and NGFC subtypes. Finally, we also observed multiple members of presynaptic GABA synthesis, release and uptake machineries including *Gad1, Syt2/10,* and *Slc6a1* differentially expressed in discrete MGE subtypes (**Figure6, Supplement1B**_i_). Collectively, these findings highlight the centrality of MGE-expressed *Grin1*-signaling during synapse formation and connectivity, which when aberrantly expressed, can lead to neurodevelopmental disorders.

## 3. DISCUSSION

### Centrality of MGE-derived interneuron-expressed NMDARs from juvenile brain

NMDARs serve as critical activity dependent signaling hubs for myriad neuronal functions due to their innate ability to directly link network dynamics to cellular calcium events, and associated transcriptional coupling. Such NMDAR-dependent excitation-transcription coupling is widely established in glutamatergic neurons [80], and in specific interneurons using candidate approaches [81] within mature circuits. However, the detailed unbiased evaluation of the transcriptional landscape of NMDAR signaling within interneurons in developing circuits undergoing refinement is lacking. Our study provides the first systematic “fingerprinting” of the transcriptional coupling associated with NMDAR signaling, exclusive to MGE-derived interneurons, providing a road map for examining NMDAR regulation of MGE-derived interneurons in a subtype specific manner.

Our unbiased transcriptional profiling approach indicates that developmental NMDAR signaling participates in MGE-derived interneuron identities by regulating the expression of transcription factors (67 genes), synaptogenic (53 genes) and connectivity factors/adhesion molecules (61 genes), and regulators of membrane excitability (78 genes), among the 802 DEGs in interneurons **(Figure7)**. We employed bioinformatic analyses to examine whether system-wide downregulation of target genes can be attributed to transcription-repression elements. Indeed, we identify a set of 10 transcriptional regulators that commonly recognize the DNA-motifs present in the identified DEGs, including the master-repressor *Rest*. Future studies are needed to examine the role of these putative repressors and and associated *cis*-regulatory elements that have not been pre-viously associated with MGE-specific transcription. However, based on broad transcriptional downregulation of target genes, we can make several predictions that should guide future investigations.

**Figure7.**
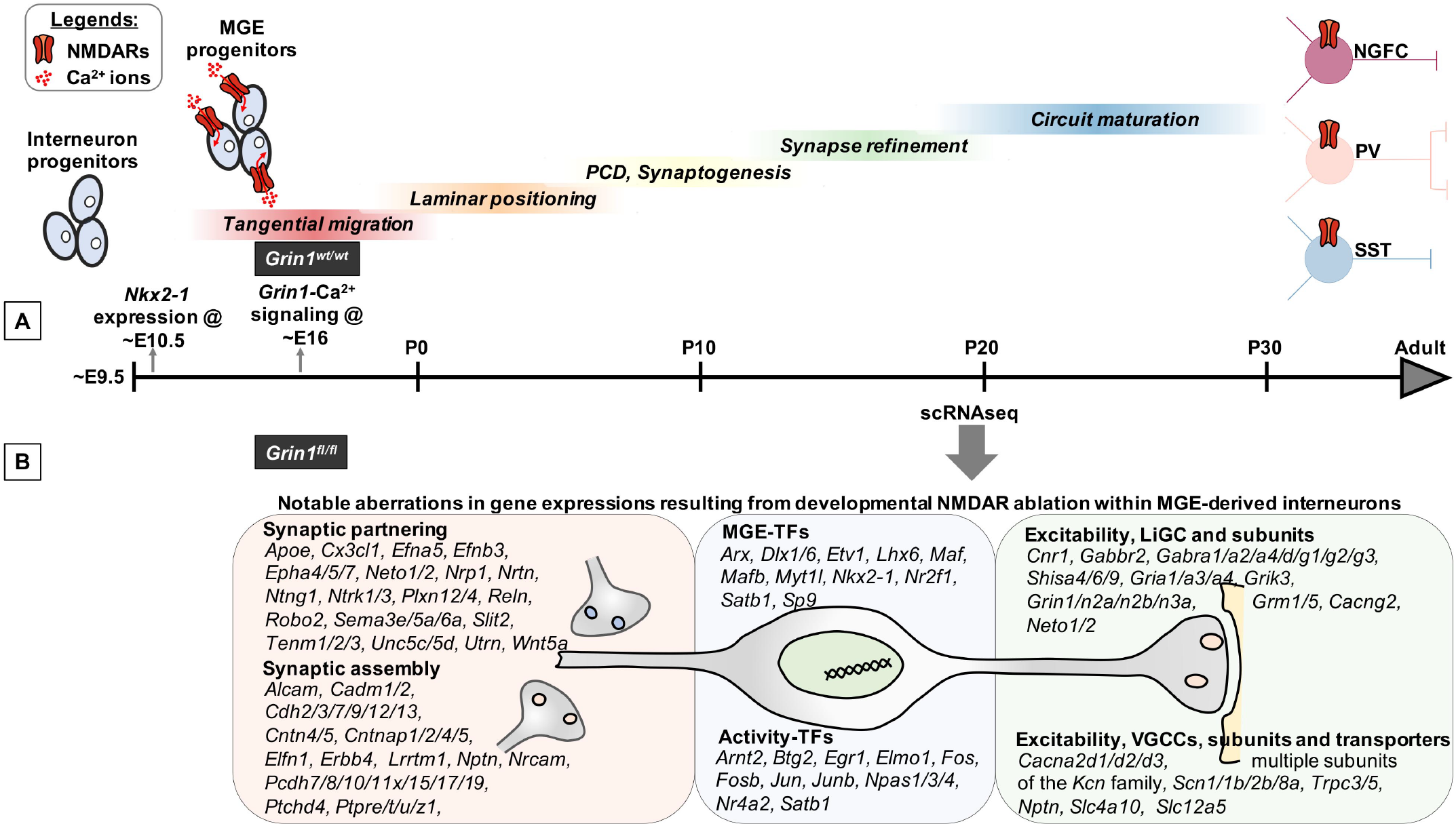
Transcriptional control of MGE development, synaptic partnering and excitablity are mediated by NMDAR signaling. **A,** *Nkx2.1* expression appears at ~ED10.5, driving MGE subtype fate in interneuronal progen-itors [42] [56]. Subsequently, their sequential developmental milestones towards circuit refinement appears to be under a combination of innate genetic mechanisms and neuronal activity. While the earliest *Grin1* expression is reported at ~ED14 in developing brain [58] [57], MGE-specific *Grin1*-mediated Ca^2+^ is recorded at ~ED16 [15]. However, the broad role played by interneuron-expressed NMDAR signaling during interneuron development until now is not well delineated. **B,** By driving *Grin1*-ablation using *Nkx2.1*-driven Cre-recombinase, we report the earliest developmental loss of NMDAR signaling, across MGE-derived interneuron subtypes. In particular, by performing scRNAseq assay in MGE-derived interneurons from the cortex and hippocampus of the mouse brain, we report a broad transcriptional aber-ration subsequent to loss of NMDAR-signaling. Notably, this expression abnormality involves numerous transcriptional factors, synaptogenic and regulators of interneuron excitability, that collectively estabish MGE subtype identities. ED, Embryonic day; PD, Postnatal day; PCD, Programmed cell death; LiGC, Ligand-gated channel; VGCC, Voltage-gated calcium channel

### Shaping interneuron identity and granularity amongst subtypes

Interneuron development from MGE is replete with combinatorial expressions of numerous transcription factors, leading to diversity [82, 83]. Several transcription factors that are impacted by NMDAR-signaling are established regulators of MGE fate, subtype abundances and identities (34 genes) including *Nkx2-1, Lhx6, Satb1, Dlx1, Dlx6, Maf, Mafb, Mef2c, Etv1, Npas1, Npas3* and *Sp9* [34, 56, 84–87] [88]. While the scRNAseq landscapes of interneurons predict several transcriptomic features that would classify them as distinct ‘cell-types’ or cell-states’ [49, 89, 90] [50] the precise mechanisms responsible for such granularity is still emerging [2].

Three key possibilities that enable NMDAR-dependent interneuron development exists, and our dataset provides potential examples for all three scenarios. (i) First, it is possible that NMDAR-dependent Ca^2+^ signaling in the developing interneuron progenitors provides a combinatorial cue that will couple with innate genetic programs to generate the diversity in interneuron subtypes. We observe key MGE-specification genes *Lhx6* and *Id2* downregulated and other genes such as *Mafb, Npas1* and *Npas3* upregulated in *Grin1*-lacking interneurons in both scRNAseq and MGE-Ribotag-seq assays. (ii) It is also possible that NMDAR signaling critically regulates neuronal activity that further shapes the expressions of MGE-subtype determining and distinct activity-dependent TFs. In line with this possibility, we establish a robust downregulation in the activity-regulated TF, *Satb1,* and a robust reduction in *Satb1*^+^ cortical interneurons that lack *Grin1*. It is notable that *Satb1* is a key activity-regulated TF that critically regulates the identity and survival of different subtypes of SST-interneurons [71, 72]. Additionally, we observe an increase in *Etv1* expression in cortical PV cells, and *Etv1* was previously demonstrated to be an activity-dependent TF that inversely correlates Ca^2+^ influx, regulating the identity of a subset of PV-interneurons [84]. (iii) Lastly, it is possible that altered neuronal activity due to aberrant NMDAR-signaling result in differential survival and/or cell-death in developing interneurons [2, 11, 21, 35]. Indicating that the *Grin1*-lacking MGE-derived interneurons have altered interneuron survival and/or cell-death, we observe differential recoveries of the subtypes within SST, PV and NGFC in the scRNAseq assay. Particularly, we observe an abundance of cortical *Chodl*-SST.1 in the *Grin1*-ablated MGE-derived interneurons, and a concomitant decrease in cortical SST.2-4 subtypes. Moreover, we report an increase in hippocampal Reln-SST2-4 subtypes upon *Grin1*-ablation by RNA *in situ* hybridization. Additionally, by independent immunostaining experiments, we observe a robust decrease in cortical MGE-derived interneurons that coexpress the TF *Satb1,* which is established to promote interneuron survival. However, detailed future studies are necessary to uncover which of the above indicated NMDAR-dependant mechanisms regulate the abundances and diversity within PV/SST/NGFC subclasses.

### Shaping interneuron subtype-specific synaptic assembly and connectivity

What is the biological context of differential expression of the TFs, in the juvenile forebrain when MGE-derived interneuron fate is assumed to be already sealed, and subtype identities established? It is emerging that some of these TFs are continually required for the maintenance of MGE fate, post development [91]. One of the ways the TFs maintain MGE subtype fate into adulthood, is by controlling the expression of genes that are essential for ongoing interneuron function. Accordingly, we predict that NMDAR-dependent ex-pression of synaptogenic and synaptic partnering molecules regulate the assembly of synapses with appropriate targets. Secreted semaphorin, ephrin, slit, netrin and neurotrophin-based signaling systems have been investigated in GABAergic neurons, during axonal pathfinding, and cell migration [92–99]. However, only recently have inroads been made into delineating their expression, and interaction with appropriate receptor systems in target synapses during accurate synaptogenesis. In addition, the NMDAR-dependent expression of synaptic adhesion molecules will further promote stability of newly formed synapses. Here, the misexpression of diverse secreted cues, their receptors and adhesion molecules by MGE subtypes during *Grin1*-ablation, provides unique insight into the molecular diversity employed during synapse establishment. Our findings also reveal numerous candidates for examining subtype specific synapse assembly, which are centrally regulated by NMDAR signaling. Of particular interest are the family of protocadherins that are reported recently to be commonly downregulated in cortical interneurons generated from Sz patient-derived induced pluripotent stem cells [100].

Subsequent to synapse formation nascent connections remain susceptible to strength modifications according to neuronal activity. Again, NMDAR-signaling in MGE-derived interneurons seems to regulate this process by the transcriptional regulation of the expressions of both presynaptic and postsynaptic members, including excitatory and inhibitory synaptic molecules and their auxiliary subunits, as well as presynaptic GABA release machinery molecules such as *Cplx1/2, Stx1b, Rab3c*. However, most dramatic is the massive down regulation of several members of the potassium channel subunits and their auxiliary subunits across MGE subtypes, with the exception of an upregulation of a few *Kcn*-genes in cortical PVs and NGFCs. While the precise impact of the diverse changes in these genes on MGE firing are currently unclear, the pattern of expression of the activity-dependent transcription factors provides us an indication.

Notable activity-dependent TFs such as *Jun, Egr1* are downregulated across all MGE subtypes, while *Fosb, Fos, Arx* are down regulated across all hippocampal MGEs, and *Satb1, Arnt2* are down-regulated across all cortical MGEs. In addition, *Npas4,* an established early-response TF [101–103] activated upon neuronal activity and Ca^2+^ influx in MGE-derived interneurons [104], was downregulated in cortical NGFCs upon *Grin1*-ablation. Lastly, *Ostn* was recently established as an activity-regulated secreted factor [105], and we observed *Ostn* to be downregulated specifically in cortical PV subtypes. Together, these changes are consistent with reduced neuronal activity in MGE subtypes upon *Grin1*-ablation, recapitulating previous reports indicating that NMDAR-antagonists can directly reduce the activity of GABAergic interneurons in adult mice [27]. Interpreting the differential expressions of activity-dependent genes during scRNAseq has been challenging, particularly, when these genes could get activated by the very process involved in cell dissociation and sorting [106, 107]. However, our use of activity-blockers and actinomycin-D throughout our MGE-*Grin1^wt^* and MGE-*Grin1^fl/fl^* scRNAseq pipelines [46] and independent validations via MGE-Ribotag-seq gives confidence that the differential expressions of activity-dependent TFs reflect biological relevance.

### NMDAR signaling in NGFCs

Among the MGE subtypes, the PV and SST interneurons are traditionally widely studied in comparison to the dendrite-targeting NGFC subtypes (that include the Ivy cells). In the present study we provide the first detailed molecular insight into the cortical and hippocampal NGFCs, subsequent to NMDAR ablation. We anticipated that these cell types could be particularly susceptible to loss of NMDARs, since we previously reported that NGFCs exhibit the most robust synaptic NMDAR conductances among the MGE subtypes [12]. Intriguingly, while the cortical NGFCs had comparable numbers of both total and unique DEGs with respect to other cortical MGE-derived interneurons **(Figure3B)**, we observed far fewer total and NGFC-specific DEGs in the hippocampus, compared to other hippocampal MGEs. However, based on the scRNAseq cell type recoveries, we predict an elaboration of NGFC.1, and a reduction in the NGFC.2 subtype upon *Grin1*-ablation. Finally, NGFCs exhibited dendritic arborization impairments subsequent to impaired NMDAR signaling [9, 11]. Indeed, we observe 49 genes among the DEGs **(Supplemental Table1)** that have established roles in regulating neuronal cytoskeleton and associated signaling, likely mediating the observed dendritic impairments in NGFCs.

### Developmental NMDAR ablation in interneurons and s chizophrenia

Impaired NMDAR function observed during human NMDAR gene mutations [108], and anti-NMDAR-encephalitis [109] results in a wide range of neuropsychiatric disorders including autism spectrum disorders [110, 111], intellectual disability [112], psychosis [113], epilepsy and associated comorbidities [114, 115]. While broadly aberrant NMDAR signaling in neurons is thought to underlie a wide range of these neurological disorders, an interneuron-centric developmental NMDAR aberration is emerging central to schizophrenia-related syndromes. Indeed, in the present study, disease mapping of the DEGs using high-ranked SZDB-Sz and SFARI-As datasets indicate that many more DEGs map with the Sz than the As database. Moreover, these disease-relevent DEGs exist in physical and functional complexes with other DEGs that are not directly mapped to the Sz database. We used only stringent, high-ranked disease genes from the database that pass several disease-relevant criteria. However, there are other DEGs that still map to lower-ranked Sz and As datasets that are ‘non-annotated’ in present study. While our study can be argued as an ‘extreme’ case of NMDAR hypofunction in MGE-derived interneurons, it provides a starting point highlighting the centrality and broad range of interneuronal NMDAR-transcriptional pathways during development.

A multitude of studies implicate NMDAR-hypofunction specific to PV cell types as a central underlying feature of schizophrenia etiology [116, 117]. However, the measurable NMDAR conductances within PV interneurons are relatively small in comparison to other MGE subtypes [12]. Additionally, NMDA signaling in non-PV interneuron subtypes drives robust dendritic inhibition in pyramidal neurons [118, 119]. Moreover, while NMDAR-ablation in *Pvalb*-Cre lines produces other behavioral deficits unrelated to the Sz-like phenotypes [30, 33], a developmental, but not adult-onset *Grin1*-ablation in *Ppp1r2*-Cre line [29] that targets a subset of PV interneurons among other subtypes [120], recapitulates core Sz-like phenotypes. Lastly, studies that map interneuron subtypes to Sz-like phenotypes indeed support the role of different interneuron classes beyond PV cells towards disease etiology [28, 33].

Integrating these ideas and based on findings from the present study, we propose the following: (i) Despite a smaller NMDAR conductance in PV interneurons, we observe a robust transcriptional coupling via NMDARs, as observed by several distinct gene expression abnormalities in this cell type relevant to human Sz. Therefore, PV-expressed NMDARs primarily serve to regulate transcriptional coupling, mediating PV-subtype abundances. (ii) The developmental window for NMDAR loss of function is particularly important because, its transcriptional regulation maintains the correct synaptogenic and assembly cues, which when lost, lead to disease-causing impaired connectivity. Perhaps, in the *Grin1^fl/fl^*: *Pvalb*-Cre mouse line, the *Grin1*-ablation occurs only at a developmental window when synaptic connectivity is sufficiently complete, explaining why the animal model does not lead to profound Sz-like impairments. (iii) The dendrite targeting SST and NGFC interneurons also exhibit robust NM-DAR signaling and transcriptional coupling. During aberrant NMDAR-transcriptional coupling, it is therefore likely that impaired dendritic connectivity and inhibition onto pyramidal neurons also contributes towards disease etiology. Therefore, our dataset provides credence to interneuronal subtype-specific granularilty, connectivity and excitability, all playing combinatorial and mutually-supporting roles during disease etiology. Taken together, our study presents a rich resource, laying the road map for systematic examination of NMDAR signaling in interneuron subtypes, by providing multiple molecular targets for examination in both normal and impaired circuits.

## 4. STAR ★ METHODS

**Table 1:**
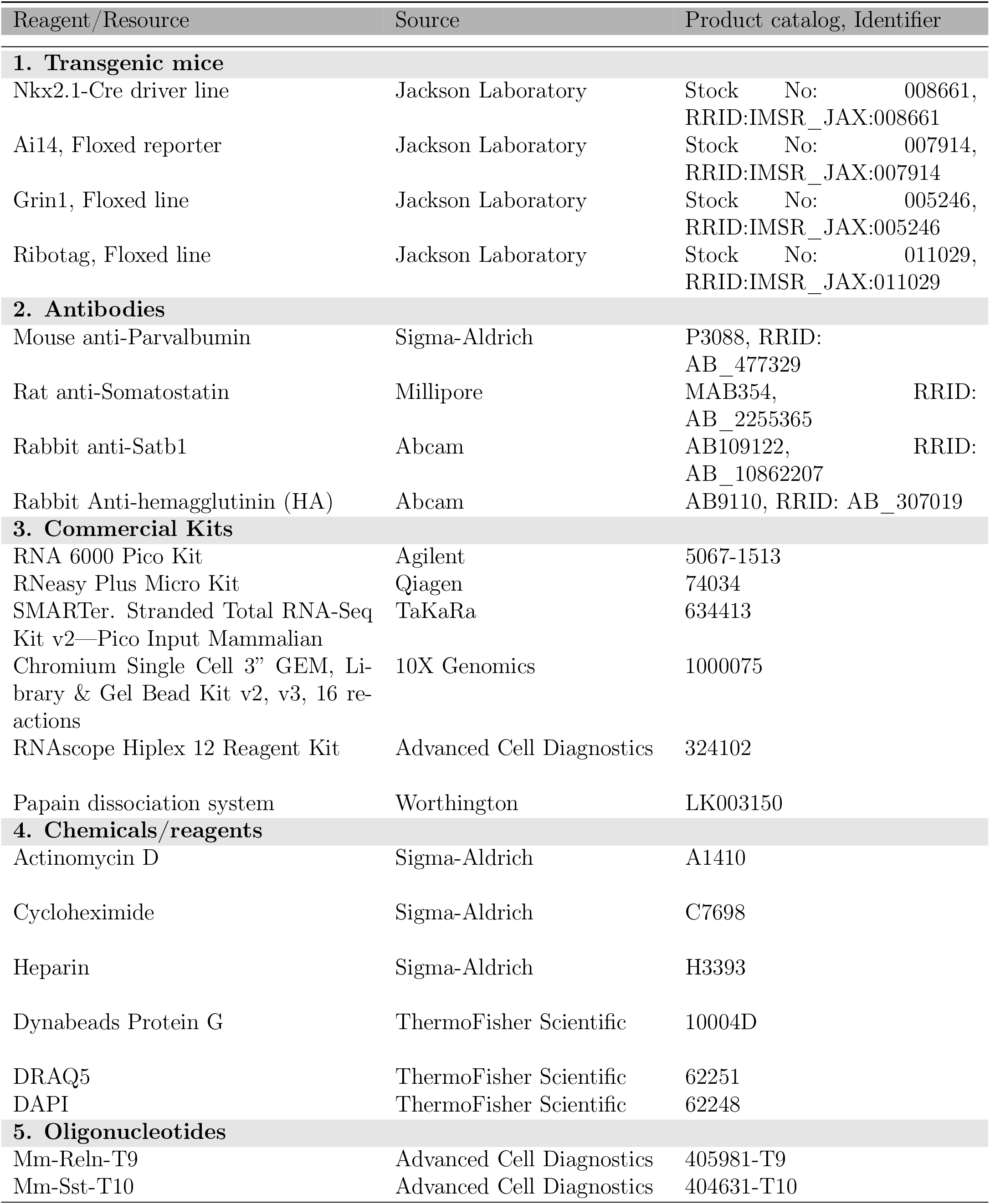
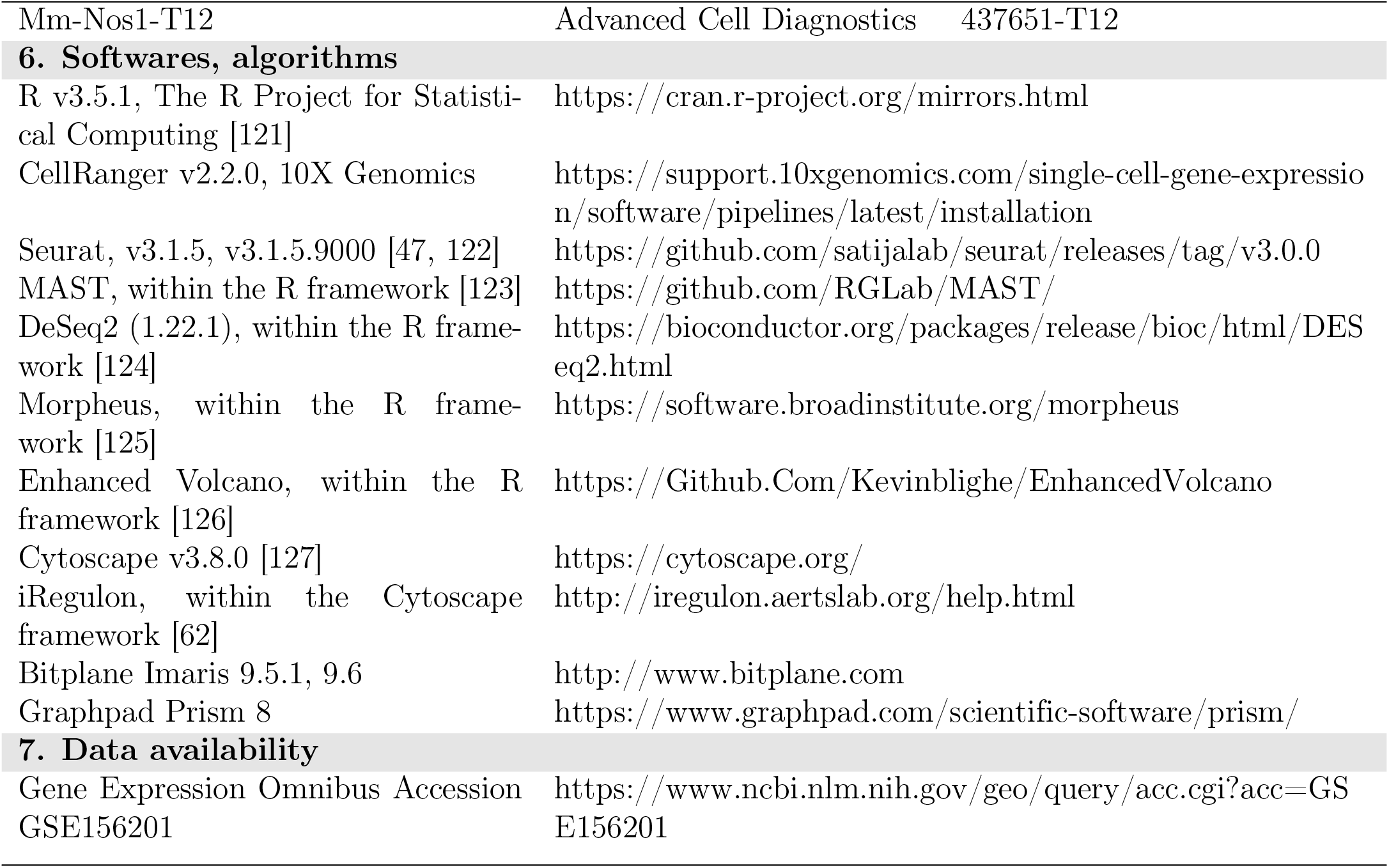
Key Resource Table

## 5. DETAILED MATERIALS AND METHODS

### Contact for Reagent and Resource Sharing

Further information and requests for resources and reagents should be directed to and reasonable requests will be fulfilled by the Lead Contact, Chris McBain.

### Animals

All experiments were conducted in accordance with animal protocols approved by the National Institutes of Health. MGE-derived interneuron-specific *Grin1^fl/fl^* line, and Ribotag crosses were conducted as indicated in Figure1-Supplement1A, and **Figure4-Supplement1D**. Littermate MGE-*Grin1^wt/wt^*controls, and MGE-*Grin1^fl/fl^* male and female mice were used during this study. Mice were housed and bred in conventional vivarium with standard laboratory chow and water in standard animal cages under a 12hr circadian cycle. Genotyping of the mice were performed as indicated in the appropriate Jackson Laboratory mice catalog.

### Single-cell dissociation and FACS

P18-20 juvenile *Nkx2*-1-Cre: Grin1^*wt/wt*^: TdT^+^ and Nkx2-1-Cre: Grin1^*fl/fl*^: TdT^+^ mice were used for single-cell sequencing experiments. All mice were anesthetized with isoflurane and then decapitated. Brain dissection, slicing and FACS sorting were carried out as described [43, 44], with slight modifications. NMDG-HEPES–based solution was used in all steps to enable better recovery of the cells [45] during FACS sorting and singlecell bar coding. Briefly, the brain sectioning solution contained NMDG-HEPES–based high-Mg^2+^ cutting solution contained 93 mM NMDG, 2.5 mM KCl, 1.2 mM NaH_2_PO_4_, 30 mM NaHCO_3_, 20 mM HEPES, 25 mM glucose, 5 mM sodium ascorbate, 3 mM sodium pyruvate, 2mM Thiourea, 10 mM MgSO4*7H2O, and 0.5 mM CaCl_2_*2H_2_O; it was adjusted to pH 7.4 with 12.1N HCl, an osmolarity of 300-310 mOsm, and carbogenated (mix of 95% O_2_ and 5% CO_2_) before use. This solution was chilled and the process of sectioning were conducted on a ice-chamber in the vibratome.

3-4, *Nkx2-1*-Cre: *Grin1^wt/wt^*: TdT^+^ or *Nkx2-1*-Cre: *Grin1^fl/fl^:* TdT^+^ mice were processed on consecutive days for single-cell sequencing experiments. TdT negative animals were processed in parallel for initially setting FACS gate for the Tomato-channel. Across the replicates, 10XMGE-Grin1-WT and 6XMGE-Grin1-null animals were used for the scRNAseq. Coronal slices containing frontal cortex and hippocampus (350mM) were cut using VT-1000S vibratome (Leica Microsystems) in cold NMDG-HEPES-based high-Mg^2+^ cutting solution. Slices were recovered in the same solution at 20°C for 30 minutes during when, they were visually inspected under fluorescence microscope and micro dissected, all under constant carbogenation. The recovery and microdissection were conducted in the NMDG-HEPES high-Mg^2+^ solution supplemented with 0.5*μ*M tetrodotoxin (TTX), 50 *μ**M*** DL-2-Amino-5-phosphonopentanoic acid (APV) and 10 *μ*M Actinomycin-D (Act-D).

Cell dissociation was performed using the Worthington Papain Dissociation System according to manufacturer instructions with minor modifications. Briefly, single-cell suspensions of the micro dissected frontal cortices or hippocampus were prepared using sufficiently carbogenated dissociation solution (containing Papain, DNAse in Earle’s Balanced Salt Solution, EBSS), supplemented with 1*μ*M TTX, 100 *μ*M APV and 20 *μ*M Act-D. After a 60 min enzymatic digestion at 37^°^C, followed by gentle manual trituration with fire-polished Pasteur pipettes, the cell dissociates were centrifuged at 300g for 5 minutes at 20^°^C, and the supernatants were discarded. The enzymatic digestion was quenched in the next step by the addition of ovomucoid protease inhibitor. Albumin density gradient was performed on the pellets, using a sufficiently carbogenated debris removal solution (containing albumin-ovomucoid inhibitor, DNAse in EBSS). The resulting cell pellets were resuspended in 1ml FACS buffer containing 10% FBS, 10U/μl of DNAse, 1*μ*M TTX, 100 *μ*M APV and 20 *μ*M Act-D in a 50:50 mix of carbogenated EBSS: NMDG-HEPES–based cutting saline (with 1mM MgSO_4_*7H_2_O, it is important to not use High-Mg^2+^ in the FACS buffer, as it interferes with the subsequent 10X scRNAseq reaction). Cells were placed in polystyrene tubes (Falcon 352235) on ice during the FACS.

For single cell sorting of TdT^+^ expressing cells by FACS, resuspended cell dissociates were filtered through 35mm cell strainer (Falcon 352235) to remove cell clumps. The single cell suspensions were then incubated with 1mg/ml DAPI (1:500) and 1mM DRAQ5 at 4°C for 5 minutes to label dead cells and live cells respectively. Samples were analyzed for TdTomato expression and sorted using a MoFlo Astrios EQ high speed cell sorter (Beckman Coulter). TdT-negative cells were used as a control to set the thresholding FACS gate for the detection and sorting of the Ai14-TdTomato-expressing cells, and the same gate was then applied for all subsequent experiments. Flow data analysis and setting of sorting gates on live (DAPI-negative, DRAQ5-positive) and Ai14-TdTomato-expressing cells were carried out using Summit software V6.3.016900 (Beckman Coulter). Per sample/session, 20,000 – 40,000 individual cells were sorted into a FBS-precoated, Eppendorf LoBind Microcentrifuge tubes containing carbogenated 10ml FACS buffer, that served as the starting material for 10X Genomics barcoding.

### 10X Genomics Chromium

The cells were inspected for viability, counted, and loaded on the 10X Genomics Chromium system, aiming to recover ~5000 cells per condition. 12 PCR cycles were conducted for cDNA amplification, and the subsequent library preparation and sequencing were carried out in accordance with the manufacturer recommendation (Chromium™ Single Cell 3’ Library & Gel Bead Kit v2 and v3, 16 reactions). Sequencing of the libraries were performed on the Illumina HiSeq2500 at the NICHD, Molecular Genomics Core facility. Replicate 1 of the scRNAseq were performed using 10X v2 reaction from which, the cell estimates, mean reads per cell (raw), median genes per cell respectively, are as follows Cortical WT: 1277, 149K, 4615; Cortical NULL: 181, 159K, 4826; Hippocampal WT: 2221, 92K, 2578; Hippocampal NULL: 404, 154K, 4903. Replicate 2 of the scRNAseq were performed using 10X v3 reaction from which, the cell estimates, mean reads per cell (raw), median genes per cell respectively, are as follows Cortical WT: 3851, 22.8K, 1536; Cortical NULL: 2898, 23.5K, 2759; Hippocampal WT: 4600, 23.6K, 850; Hippocampal NULL: 4436, 25.8K, 3143. Replicate 3 of the scRNAseq were performed using 10X v3 reaction from which, cell estimates, mean reads per cell (raw), median genes per cell respectively, are as follows Cortical WT: 3960, 24.8K, 2870; Hippocampal WT: 3159, 26.9K, 2956. Representative quality metrics from Replicate 2 are indicated in **Figure1-Supplement1B,C,D,E.** Demultiplexed samples were aligned to the mouse reference genome (mm10). The end definitions of genes were extended 4k bp downstream (or halfway to the next feature if closer), and converted to mRNA counts using the Cell Ranger Version 2.1.1, provided by the manufacturer.

### MGE-Ribotag-seq

This assay was performed as recently described [70]. Notably, *Nkx2.1^cre^: Grin1^fl/wt^* were bred with Ribotag mice for >10 generations to obtain homozygosity in *Rpl22^HA/HA^.* Age-matched *Nkx2.1^cre^*: *Grin1^wt/wt^*: *Rpl22^HA/HA^* and *Nkx2.l^cre^*: *Grin1^fl/fl^*: *Rpl22^HA/HA^* littermates (2 males and 2 females per genotype) were utilized for the MGE-Ribotag-seq. RNAs bound with anti-HA immunoprecipitates and RNA from bulk tissue (input) were purified using RNeasy Plus Micro Kit and the quality of RNA were measured using RNA 6000 Pico kit and 2100 Bioanalyzer system. cDNA libraries were constructed from 250 pg RNA using the SMARTer Stranded Total RNA-Seq Kit v2 only from samples with RNA Integrity Numbers > 6. Sequencing of the libraries were performed on the Illumina HiSeq2500, at 50 million 2 x 100bp paired-end reads per sample. ~75% of reads were uniquely mapped to genomic features in the reference genome. Bioconductor package DESeq2(1.22.1) was used to identify differentially expressed genes (DEG). This package allows for statistical determination of DEGs using a negative binomial distribution model. The resulting values were then adjusted using the Benjamini and Hochberg’s method for controlling the false discovery rate (FDR).

### Data processing, analyses, visualization and differential expression testing

Processing (load, align, merge, cluster, differential expression testing) and visualization of the scRNAseq datasets were performed with the R statistical programming environment [128] (v3.5.1) and Seurat package (v3.1.5, a development version of Seurat v3.1.5.9000 was used to generate violin plots in 2C and 5B) [47, 48]. Data set preprocessing, comparison of WT- and NULL-Ai14 cells, canonical correlation analyses, and differential expression of genes (p_*adj*_ < 0.01) within the same cluster between WT- and NULL-Ai14 cells were performed according to default Seurat parameters, unless otherwise mentioned. Quality control filtering was performed by only including cells that had between 200-6000 unique genes, and that had <30% of reads from mitochondrial genes. While the WT replicates had no cells above 30% mitochondrial genes, only NULL replicates from both brain regions exhibited 7-12% of cells above this threshold. Suggestive of inherent biological impact of *Grin1*-ablation, we repeated the clustering and subsequent analyses without excluding any cells. These analyses did not alter the clustering or skew the gene list. Clustering was performed on the top 25 PCs using the function FindClusters() by applying the shared nearest neighbor modularity optimization with varying clustering resolution. A cluster resolution of 1.0 was determined to be biologically meaningful, that yielded all known MGE cardinal classes. Initial analyses were performed on the WT datasets separately (WT.alone), and similar set of analysis parameters were applied when the WT and NULL samples were merged (WT.NULL.integrated) for subsequent differential expression testing. Phylogenetic tree relating the ’average’ cell from each identity class based on a distance matrix constructed in gene expression space using the BuildClusterTree() function. Overall, we identified 27, and 33 clusters using this approach in the WT.alone, and WT.NULL.integrated assays respectively. The WT.alone correspond to 11 MGE.GAD1/2 clusters **(Figure1&2)**, while the WT.NULL.integrated assay correspond to 12 clusters **(Figure5-Supplement.1).** We first searched for the top differential markers for each MGE subcluster using the FindAllMarkers() function. The genes thus identified for the integrated data is presented in ***Supplemental Table1b.*** Determination of MGE and non-MGE identities are performed based on existing interneuron literature and other scRNAseq datasets [1, 43, 50, 53, 89, 91, 129–131]. The labels from Figures 1 and 2 are matched with the top gene markers identified by the FindAllMarkers() function and the similarly named clusters in Figures 1 and 2 have the same identities. Lastly, for the integrated analyses and differential expression testing, we first merged the identities of the subclusters SST.1-SST.6 and TH.1, and relabelled as SST subset; PVALB.1-3 relabelled as PVALB subset; and NGFC.1-2 relabelled as the NGFC subset during subsequent analysis **(Figure3)**.

Differential gene expression testing were performed using the MAST package within the FindMarkers function to identify the differentially expressed genes between two subclusters. MAST utilizes a hurdle model with normalized UMI as covariate to generate the differential fold changes [123], and is known to result in underestimation of the magnitude of fold change (FC) [132]. Therefore, while applying a stringent false-discovery rate <0.01, we determined the minimum FC based on the control gene *Grin1*, which is the target gene knocked out in MGE-derived interneuron celltypes. Notably for *Grin1*, we had previously demonstrated that the NGFCs which carry maximum NMDAR component among MGEs, are devoid of NMDAR current at this comparable age [9]. In the present scRNAseq assay, we observe a logFC for *Grin1* ranging between −0.1 to −0.35 across both brain regions and all MGE subtypes. Therefore, we determined a minimum logFC in our DEGs as ±0.1 to be meaningful. Previous studies have demonstrated the MAST approach for DEG testing to be powerful in determining subtle changes in highly transcribed genes, and among abundant populations, additional to under representing changes among weakly transcribed genes [123, 132]. Volcano plots and Heat maps for the DEG were generated using EnhancedVolcano package and Morpheus package within the R framework. Associated scRNAseq source data, including sequencing reads and single cell expression matrices, is available from the Gene Expression Omnibus (GEO) under accession number GSE156201.

### Pathway analyses PPI network mapping and disease mapping

Ingenuity Pathway Analyses were conducted on the differentially expressed genes to generate the molecular functional annotation and to identify the biological pathways and disease pathways overrepresented. This tool was also used to annotate genes with their known cellular functional classes. Additional Gene Ontology mapping and KEGG analyses were conducted using ShinyGO [133]. Protein-protein interaction (PPI) mapping datasets from a variety of curated databases [134–136] were conducted as previously described [76] [137] using the Cytoscape [127] platform (v3.8.0). Schizophrenia risk genes integrated from various sources including genomewide association studies (GWAS), copy number variation (CNV), association and linkage studies, post-mortem human brain gene expres-sion, expression quantitative trait loci (eQTL) and encyclopedia of DNA elements (ENCODE), were downloaded from http://www.szdb.org/ [73]. Autism Spectrum Disorder risk genes integrated from various sources were downloaded from Simons Foundation https://gene.sfari.org/ [74]. SZDB genes that had a integrated total score of 3-6 (1419 genes, 22% out of 6387) were considered ‘high-risk’ for DEG mapping **(Supplemental Table5a)**. SFARI genes scored 1-2 with accounting for a high strength of evidence (392 genes, 42% out of 943), were considered ‘high-risk’ for DEG mapping **(Supplemental Table5b)**. Transcriptional factor motif enrichment search using the iRegulon was also conducted using Cytoscape using default parameters.

### Immunostaining

All solutions were freshly prepared and filtered using 0.22*μ*M syringe filters for parallel treatments of wildtype and MGE-Grin1-null groups. Adult mice of postnatal day (PD) 30/60/210 were Mice were deeply anesthetized with isoflurane and perfused transcardially with 1X phosphate buffer saline (PBS) and followed by the fixative 4% paraformaldehyde. The brains were post-fixed in the same fixative for overnight at 4^°^C for the immunostaining assays. Postfixed brains were serially dehydrated using 10%/20%/30% sucrose solutions at 4 °C. Coronal sections (50 *μ*M) were cut on a freezing microtome. Immunostaining was performed on free-floating sections. Tissue sections were permeabilized and blocked in 1 × PBS + 1% bovine serum albumin + 10% normal goat serum + 0.5% Triton X-100 (Carrier PB) at room temperature for 2 h, followed by incubation in primary antibodies, listed below, diluted with 1 × PBS + 1% bovine serum albumin +1% normal goat serum + 0.1% Triton X-100 overnight at 4^°^C. Tissue sections were then incubated with secondary antibodies, listed below, diluted in Car-rier Solution (1:1000), and DAPI (1:2000) at room temperature for 1–2 h and mounted on Superfrost glass slides, and coverslipped using Mowiol mounting medium and 1.5 mm cover glasses.

### Antibodies

The following primary antibodies were used: mouse anti-PV (1:1000; Sigma-Aldrich Cat# P3088, RRID: AB_477329), rat anti-SST (1:1000; Millipore Cat# MAB354, RRID: AB_2255365), rabbit anti-Satb1 (1:1000; Abcam Cat# ab109122, RRID: AB_10862207), rabbit anti-HA (Abcam, Cat#ab9110, RRID AB_307019). Secondary antibodies were conjugated with Alexa Fluor dyes 488 or 633 (1:1000; Thermo Fisher Scientific).

### RNAscope HiPlex *in situ* hybridization

All apparatus, dissection instruments for this assay were treated to maintain RNAse-free conditions. Brains from two sets of P20-25 littermates from both genotypes were rapidly dissected and rinsed in fresh ice-cold RNAse-free 1XPBS. Individual brains were placed into cryomolds and snap-freezed by dipping into isopentene:dry ice mix. Frozen brains were stored in −80°C. 15*μ*M fresh frozen saggital sections were obtained using a RNAse-free cryostat, and sections serially mounted on RNAse-free Superfrost glass slides. After a series of pretreatments with 4%fresh PFA and dehydration using 50%, 70% and 100% EtOH, the slides were processed for HiPlex assay according to ACDBio manufacturer’s instructions within a week. A panel of HiPlex probes were utilized counterstained with DAPI (1:2000), but only data from *Reln*, *Sst*, *Nos1* are presented in this study.

### Image acquisition and analysis

#### Immunostaining

Mouse brains from 3-6 ani-mals (2-3 sections per animal) were used for each condition, and section depth were matched between the genotypes for parallel immunostaining. Fluorescent images were captured using the 20X objective of a Nikon Widefield Fluorescence, Spinning Disk Confocal microscope. For all slices with immunostained or genetically reported signal, 50 *μ*M thin sections were imaged using 10x/0.45 CFI PlanApo objective (imaging settings: Numerical Aperture 0.75, bit depth 16-bit, Exposure 100ms). Confocal stacks were stitched using NIS Elements (Nikon) before importing them into Imaris software (Bitplane, version 9.6). Cell bodies were marked in Imaris software using the ‘Spots’ function. *Nkx2-1*-Cre:TdT^+^ RFP+, PV+, SST+, SATB1+ cell bodies were detected using the automatic function, with a signal detection radius of 15 *μ*M. The Imaris ‘Quality’ filter was set above an empirically determined threshold to maximize the number of detected cells while minimizing observed false positives. SST+ cell bodies were marked manually using the Imaris ‘Spots’ function. ROI 3D borders around hippocampus or somatosensory cortex, drawn manually using the Imaris function ‘Surfaces’. Spots were then split within each ROI using the Imaris function ‘Split Spots’. Overlap of RFP+ cells with other markers (PV, SST, SATB1) was addressed by filtering the RFP+ Spots above an empirically determined threshold intensity in the channel relative to the marker of interest. Each image with an automatic analysis by Imaris was checked by an expert and incorrectly identified cell bodies where refined if required. Total numbers of cell counts are normalized to the volume of the section imaged, and normalized to the Ai14 cell numbers where applicable. In **Figure4E, Figure2-Supplement1A,B** error bars reflect standard error of mean (s.e.m); Two-tailed unpaired t-test was performed using Prism8.

#### RNAscope-HiPlex

Mouse brains from 2 animals (2-3 sections per animal) were used for each condition, and section depth were matched between the genotypes for parallel HiPlex assay. Fluorescent images were captured using the 40X-oil objective of a Nikon Widefield Fluorescence, Spinning Disk Confocal microscope. All sections were imaged using 10x/0.45 CFI PlanApo objective (imaging settings: Numerical Aperture 0.75, bit depth 16-bit, Exposure 100ms). Confocal stacks were stitched using NIS Elements (Nikon) before importing them into Imaris software (Bitplane, version 9.6). *Sst* + *Reln**+** Nos1* + RNA signals that contain DAPI were identified using ‘Spot’ function as described above, and the overlap of *Sst* + spots with *Reln**+*** or *Nos1* + spots were identified by thresholding the signals of *Reln* or *Nos1* respectively. Total numbers of cell counts are normalized to the area of the thin section imaged. In **Figure2-Supplement1C,D** error bars reflect standard error of mean (s.e.m); Two-tailed unpaired t-test was performed using Prism8.

## 6. AUTHOR CONTRIBUTIONS

VM and CJM conceived the project. VM, TJP and CJM designed the experiments, DM performed FACS sorting and analysis. VM, YZ, TJP performed 10X scRNAseq. VM performed Ribotag assays. VM, AM, CJR, CE, RD performed 10X scRNAseq bioinformatic analyses. VM, XY and AP conducted RNAscope, immunuflourescent staining, imaging and analysis. CJM supervised the study. VM and CJM wrote the paper and all authors edited the manuscript.

## ACKNOWLEDGEMENT

This work was supported by *Eunice Kennedy Shriver* NICHD Intramural Award to CJM. We thank Steven L. Coon, Tianwei Li and James R. Iben at the Molecular Genomics Core, NICHD, for RNA sequencing and bioinformatics support. We thank Vincent Schram and Lynne Holtzclaw of the NICHD Microscopy and Imaging Core for imaging support, and we thank Carolina Bengtsson Gonzales for assistance with improving cell viability during dissociation and FACS. We also thank Steven Hunt, Daniel Abebe for assistance with animal colony maintenance.

---

**Figure1, Supp.1.**
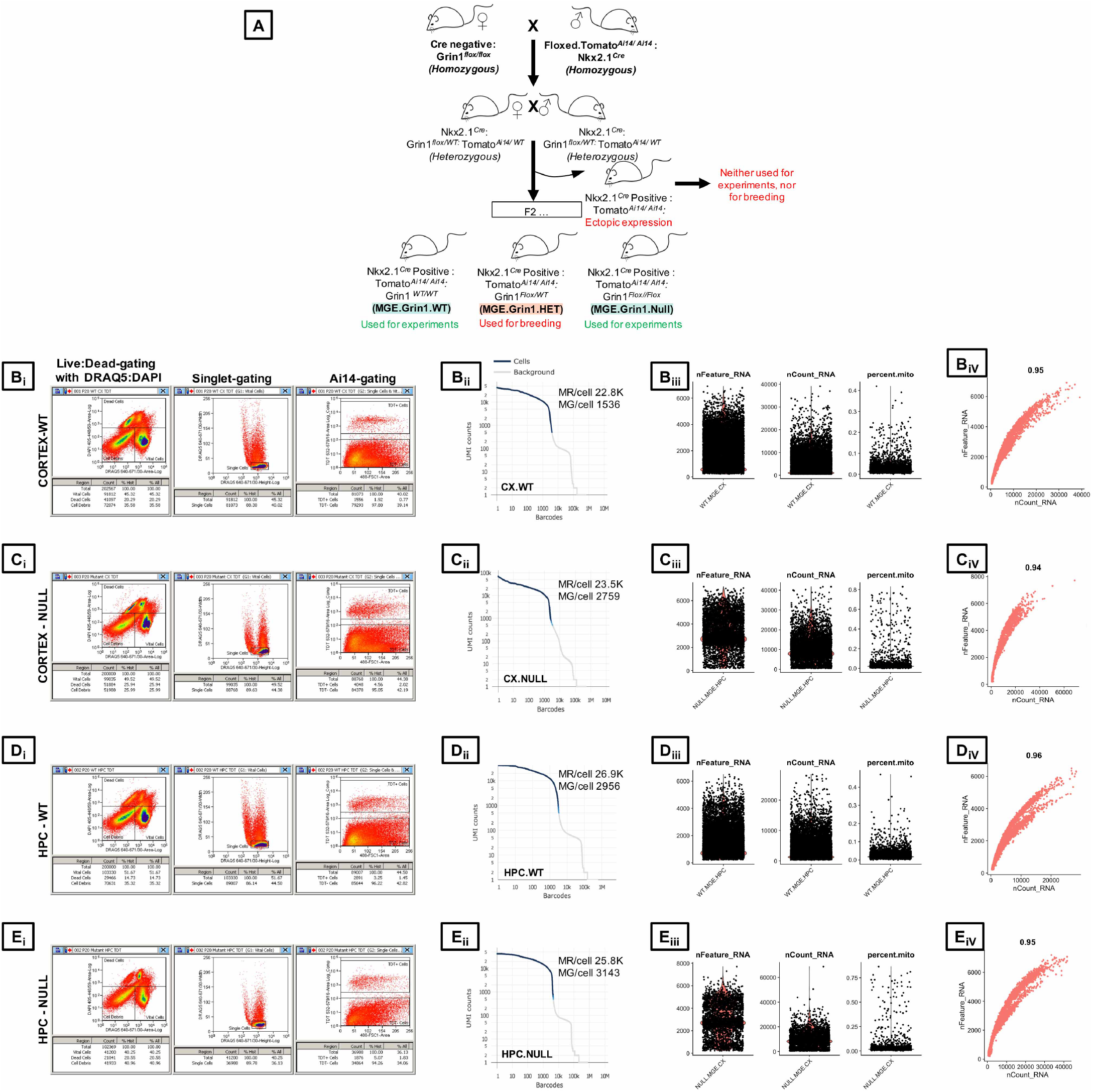
Schematic overview and quality control for scRNAseq. **A,** Breeding strat-egy, **B_i_,C_i_,D_i_,E_i_**, Representative FACS gates to sequentially isolate live: dead cells using DAPI: DRAQ5 staining, singlet-gating and TdT^+^-reporter gating to obtain reporter-positive MGE-derived interneurons from frontal cortex and hippocampus. **B_ii_,C_ii_,D_ii_,E_ii_**, Barcode Rank Plots for cells from WT and NULL mice, demonstrating separation of cell-associated barcodes and those associated with empty partitions. UMI, unique molecular identifier; MR, Mean Reads; MG, Median Genes. **B_iii_,C_iii_,D_iii_E_iii_**, Distributions of the total number of genes, percentage of mitochondrial genes and UMIs per cell in control mice **B_iv_,C_iv_,D_iv_,E_iv_**, Pearson correlation coefficient of the distributions of the total number of genes and the UMI

**Figure1, Supp.2.**
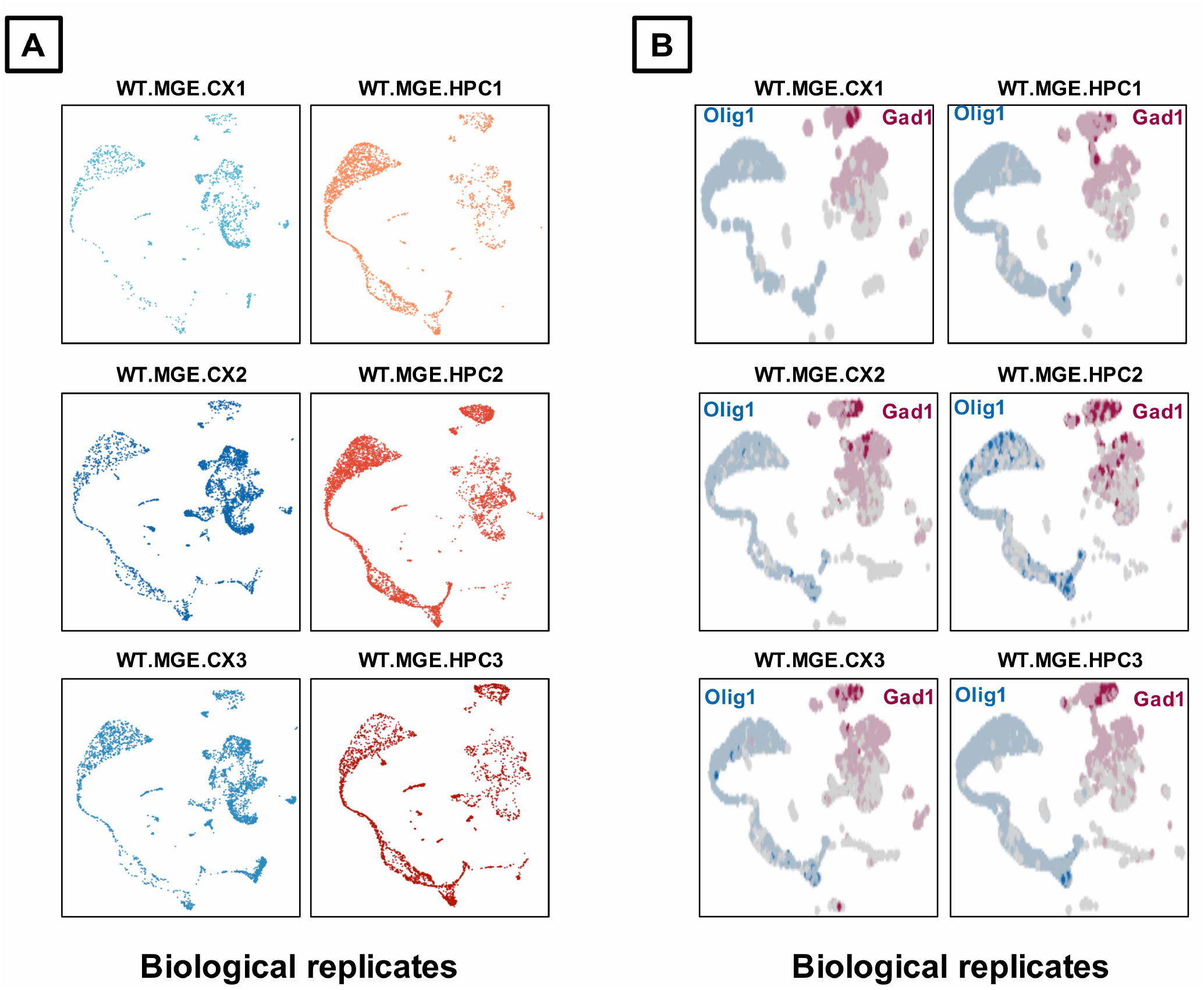
Biological replicates. Representative UMAP plots of the 3 biological replicates from cortex and hippocampus indicates **A,** similar clustering, and **B,** similar expression profiles of *Nkx2-1*-derived, *Gad1*-expressing MGE-derived interneurons and *Nkx2-1* derived, *Olig1*-expressing oligodendrocytes.

**Figure1, Supp.3.**
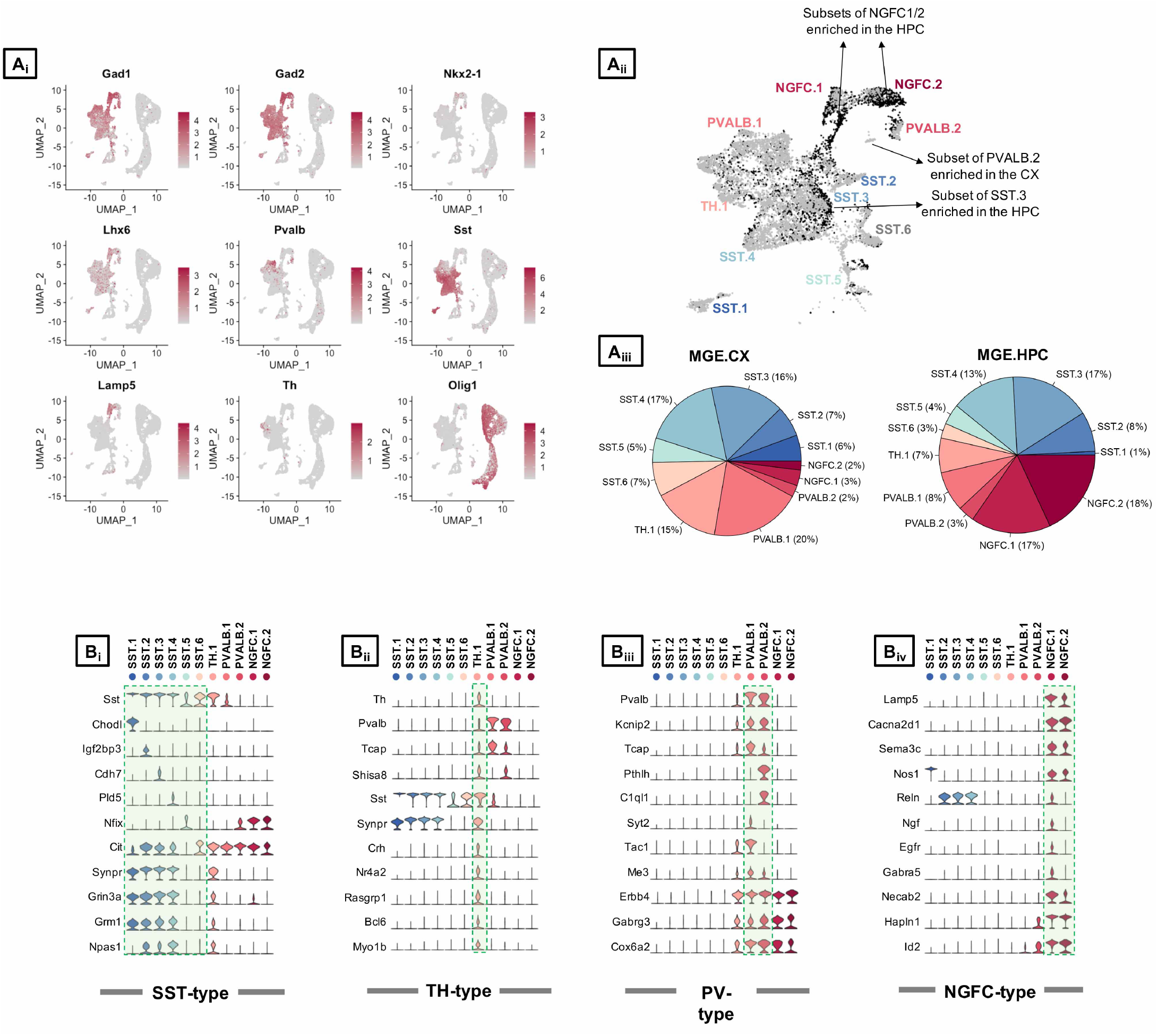
Select marker gene expression across the subtypes of merged cortical and hippocampal MGE-*Grin1^wt/wt^*. **A**_i_, UMAP representation of cardinal MGE markers genes in the cortical and hippocampal merged dataset. **A**_ii_, UMAP representation colored by region, highlighting the region-specific enrichments of MGE subsets. **A**_iii_, Pie chart indicating the percentages of cells recovered across the interneuron subtypes from cortex and the hippocampus. **B,** Violin plot showing the distribution of expression levels of well-known representative cell-type-enriched marker genes across the 11 MGE subtypes.

**Figure1, Supp.4.**
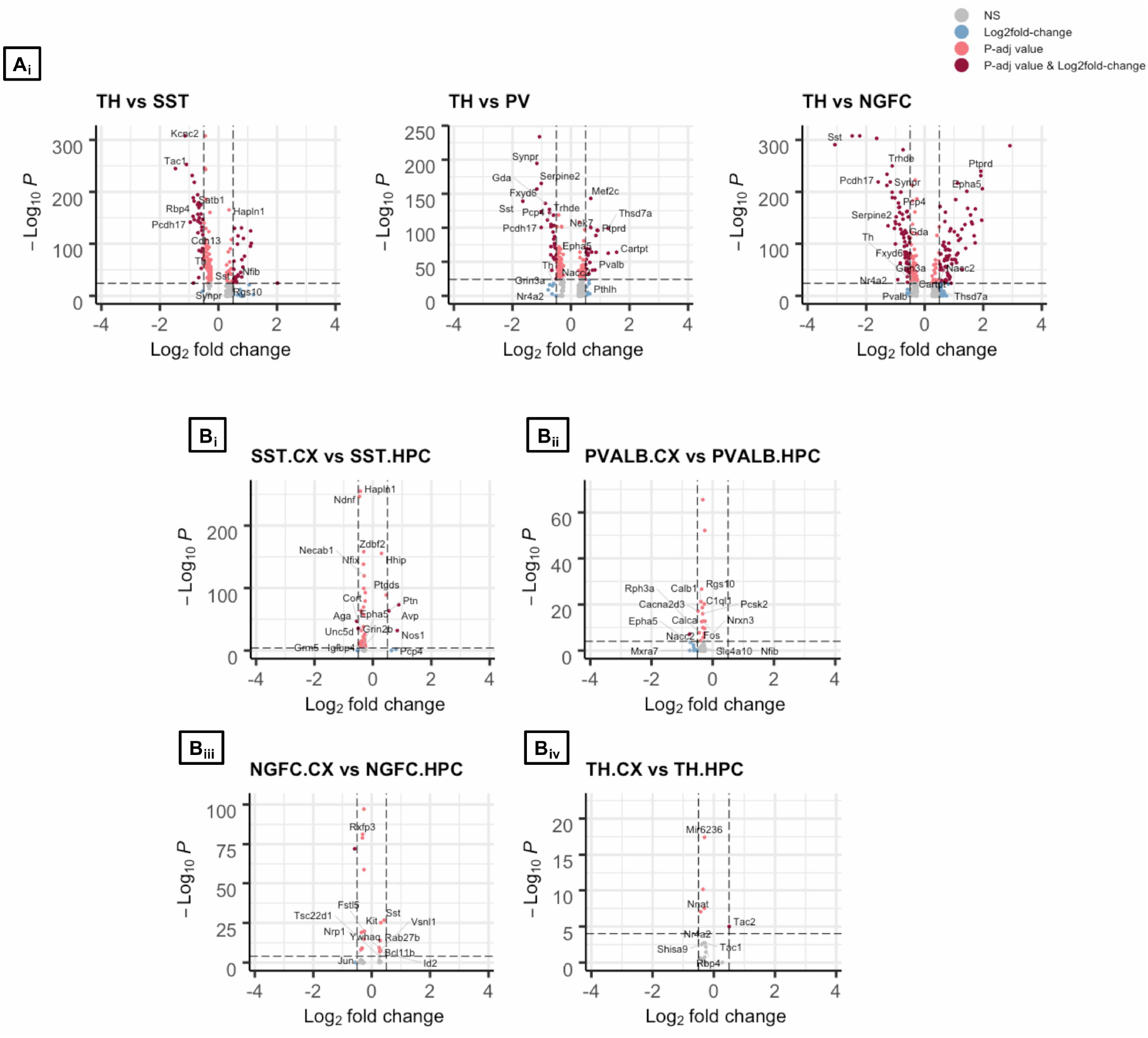
MGE subtype differences between cortex and hippocampus. Volcano plot representing the –log10 False Discovery Rate (FDR) versus log2 fold change (FC) between **A,** TH-expressing MGE subsets and the remaining MGE subset SST, PVALB and NGFC; **B,** Differential expression of the cardinal MGE classes between cortex and hippocampus, at a fold change ≥0.5 and FDR <10e-5.

**Figure1, Supp.5.**
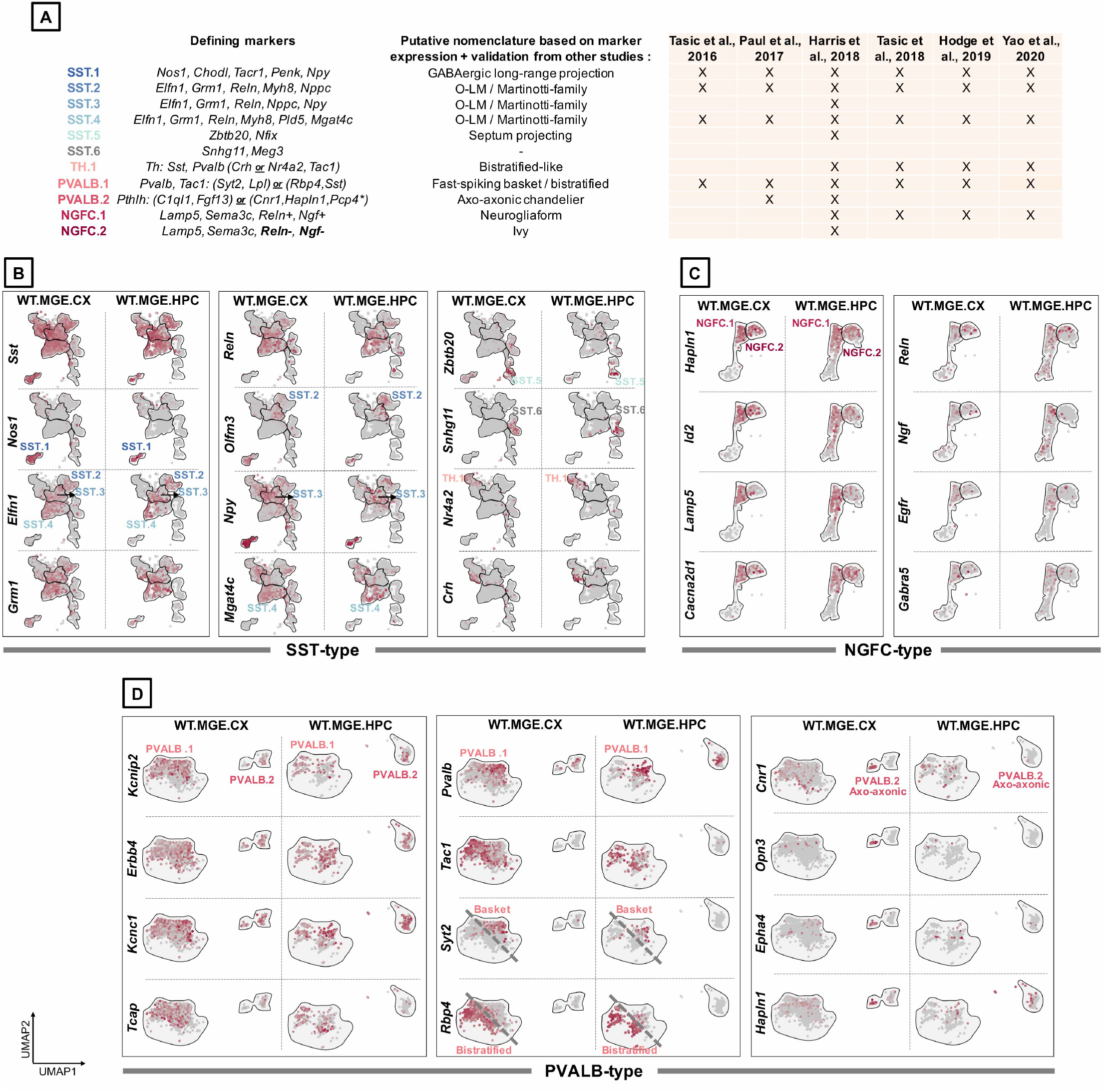
MGE subtype annotation based on marker expression. **A,** Table indicating the subype-defining marker genes observed in the present study and their descriptions in the previous scRNAseq datasets (*indicates the genes expressed in the cortex-exclusive PVALB.2 subcluster). Representative UMAP plots of MGE subtype-enriched genes in **B,** SST subclusters, **C,** NGFC subclusters, and **D,** PVALB subclusters.

**Figure2, Supp.1.**
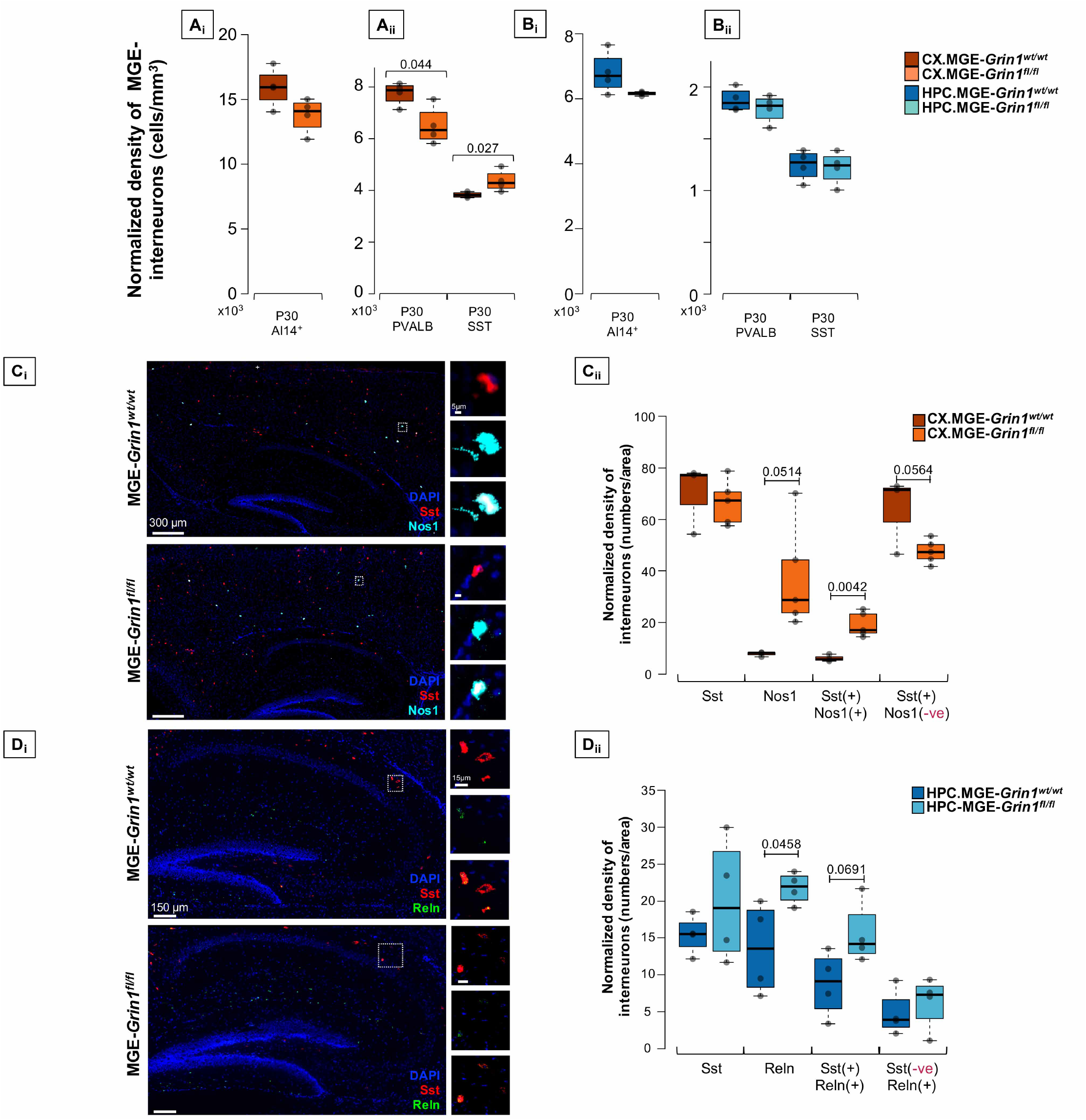
Validation of MGE subtype abundances by immunostaining and RNA *in situ* hybridization subsequent to *Grin1*-ablation. **A,** Boxplots indicating the cell counts of MGE-derived interneurons expressing **(A_i_**) Ai14/tdTomato, (**A_ii_**) PV, SST immunostaining from P30 somatosensory cortex of *Grin1*^wt/wt^and *Grin1* fl/fl. **B,** Boxplots indicating the cell counts of hippocampal MGE-derived interneurons expressing (**B_i_**) Ai14/tdTomato, and (**B_ii_**) PV, SST immunostaining from P30 *Grin1^wt/wt^* and *Grin1^fl/fl^.* **C,** Exami-nation of cortical Nos1-expressing Chodl-SST.1 subtype abundances by (**C_i_**) in situ hybridization using *Sst* and *Nos1* RNAscope probes from P20-25 somatosensory cortex, counterstained with DAPI. (**C_ii_**) Boxplots indicate Sst^(+)^, Nos1^(+)^, SST^(+)^:Nos1^(+)^or SST^(+)^:Nos1^(-)^ cell counts.**D,** Examination of hippocampal Reln-expressing SST.2-4 subtype abun-dances by (**D_i_**) in situ hybridization using *Sst* and *Reln* RNAscope probes from P20-25 hippocampus, counterstained with DAPI. (**D_ii_**) Boxplots indicate Sst^(+)^, Reln^(+)^, SST^(+)^:Reln^(+)^or SST^(-)^:Reln^(+)^ cell counts. n = 4-6 brains from each genotype for immunostaining; n = 2 brains (4-6 sections) from each genotype for RNAscope. Error bars reflect s.e.m.; two-tailed unpaired *t*-test, for statistical analysis.

**Figure2, Supp.2.**
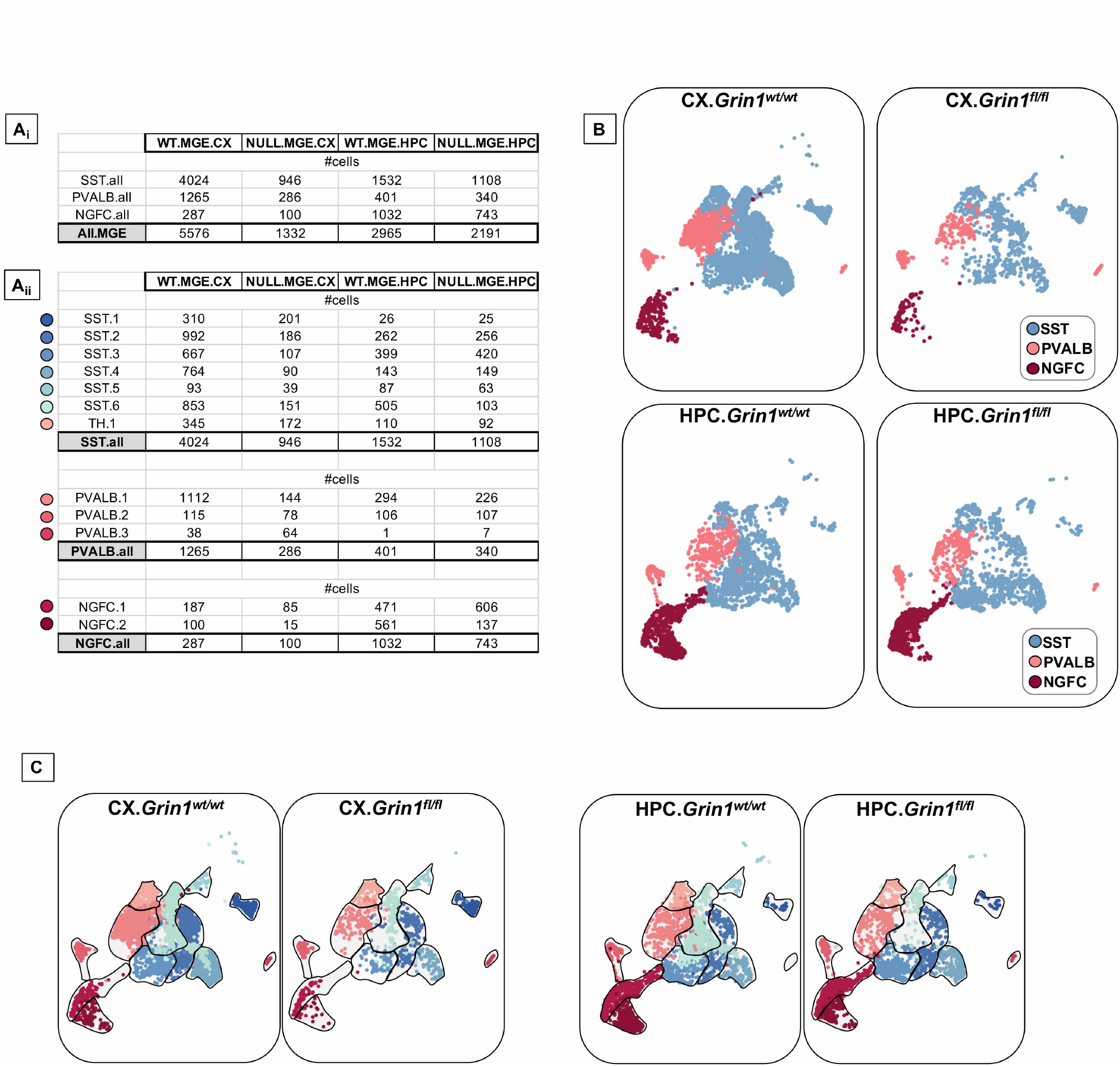
scRNAseq differential recoveries of MGE subtypes. **A**_i_, Number of cells recovered across cardinal subtypes SST, PV and NGFC. **A**_ii_, Number of cells recovered within the subtypes of PV / SST/ NGFC. **B,** UMAP representation colored by cardinal MGE-derived interneuron subtypes SST, PVALB and NGFC, highlighting the differential enrichments of cells **C,** Representative UMAP plots indicating the granularity among PV/SST/NGFC subtypes between both brain regions and both genotypes.

**Figure2, Supp.3.**
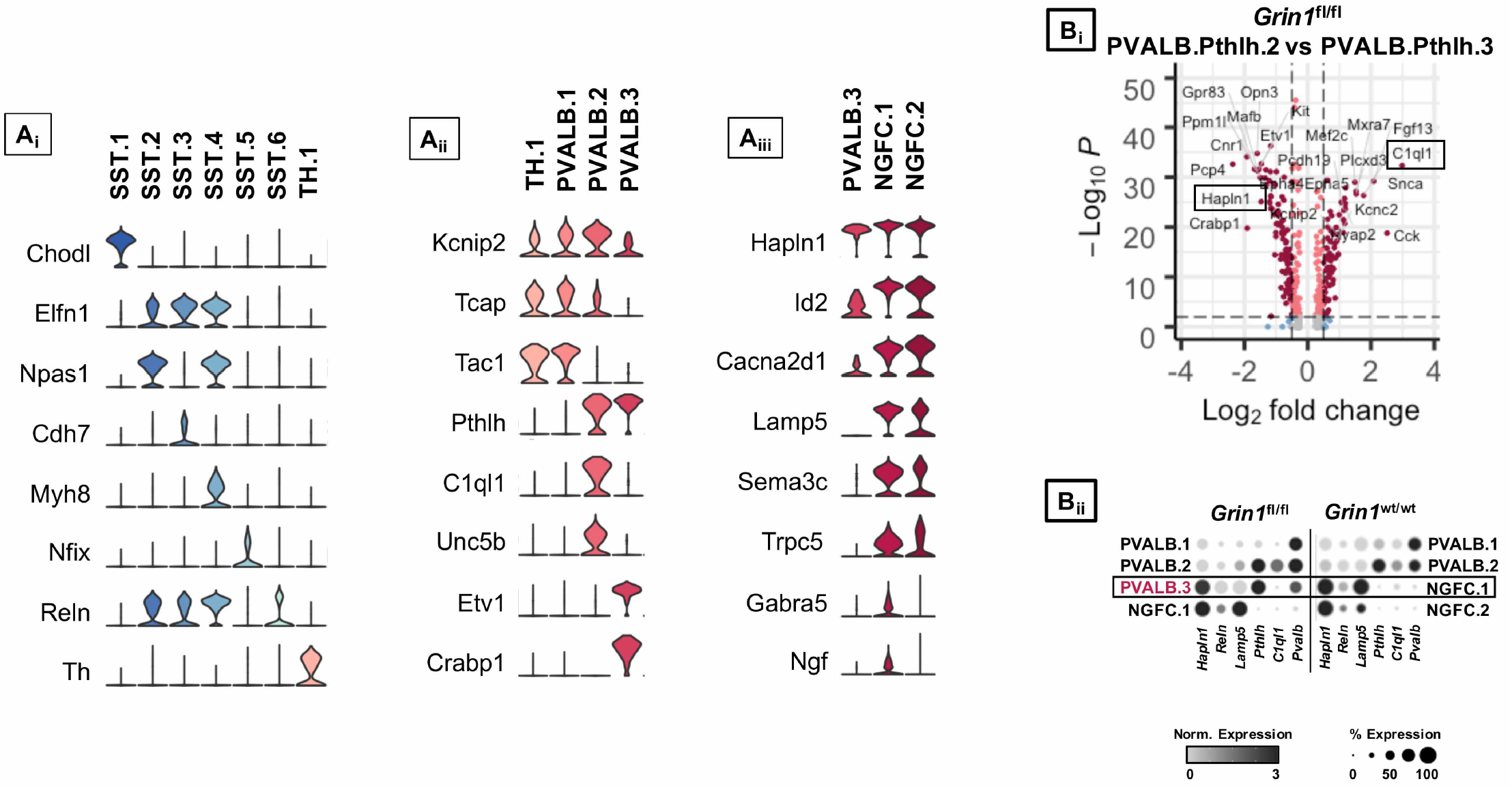
Select marker gene expression across the subtypes of merged MGE-derived interneurons from *Grin1^wt/wt^* and *Grin1^fl/fl^*. **A,** Violin plot showing the distribution of expression levels of well-known representative cell-type-enriched marker genes across the MGE subtypes. **B**_i_, –log 10 False Discovery Rate (FDR) versus log2 fold change (FC) between *Pthlh-PVALB.2* and *Pthlh*-PVALB.3 at a fold change ≥0.5 and FDR <10e-3. **B**_ii_, Dot plots representing the normalized expressions of NGFC marker genes mis expressed in *Pthlh*-PVALB.3 upon *Grin1*-ablation.

**Figure2, Supp.4.**
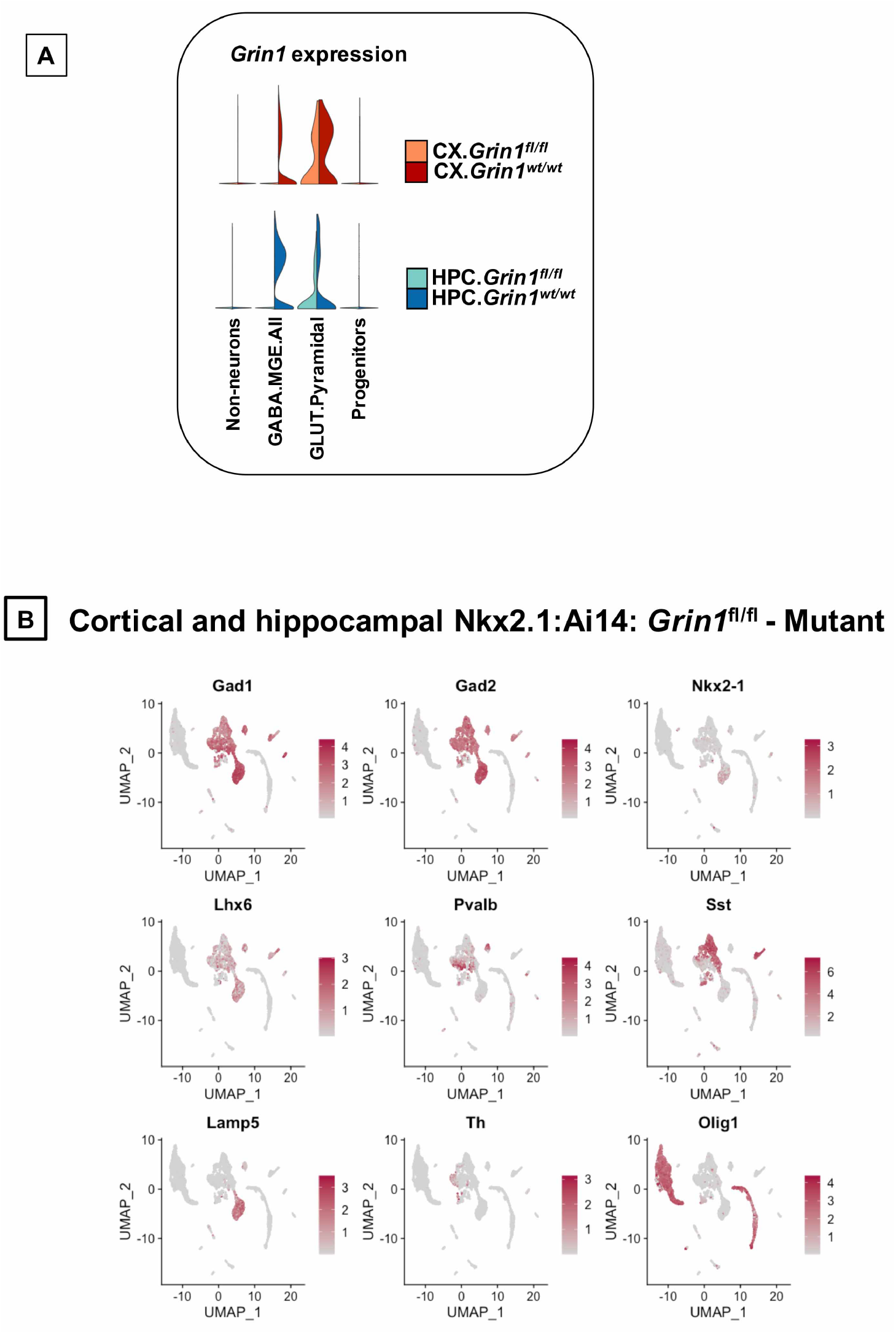
Expression of control genes in the MGE subtypes subsequent to *Grin1*-ablation. **A**_i_, Split-violin plot from both genotypes indicating the expression of *Grin1* in the MGE-derived interneurons, pyramidal neurons and non-neurons. **B,** Representative UMAP plots of cardinal MGE markers genes.

**Figure3, Supp.1.**
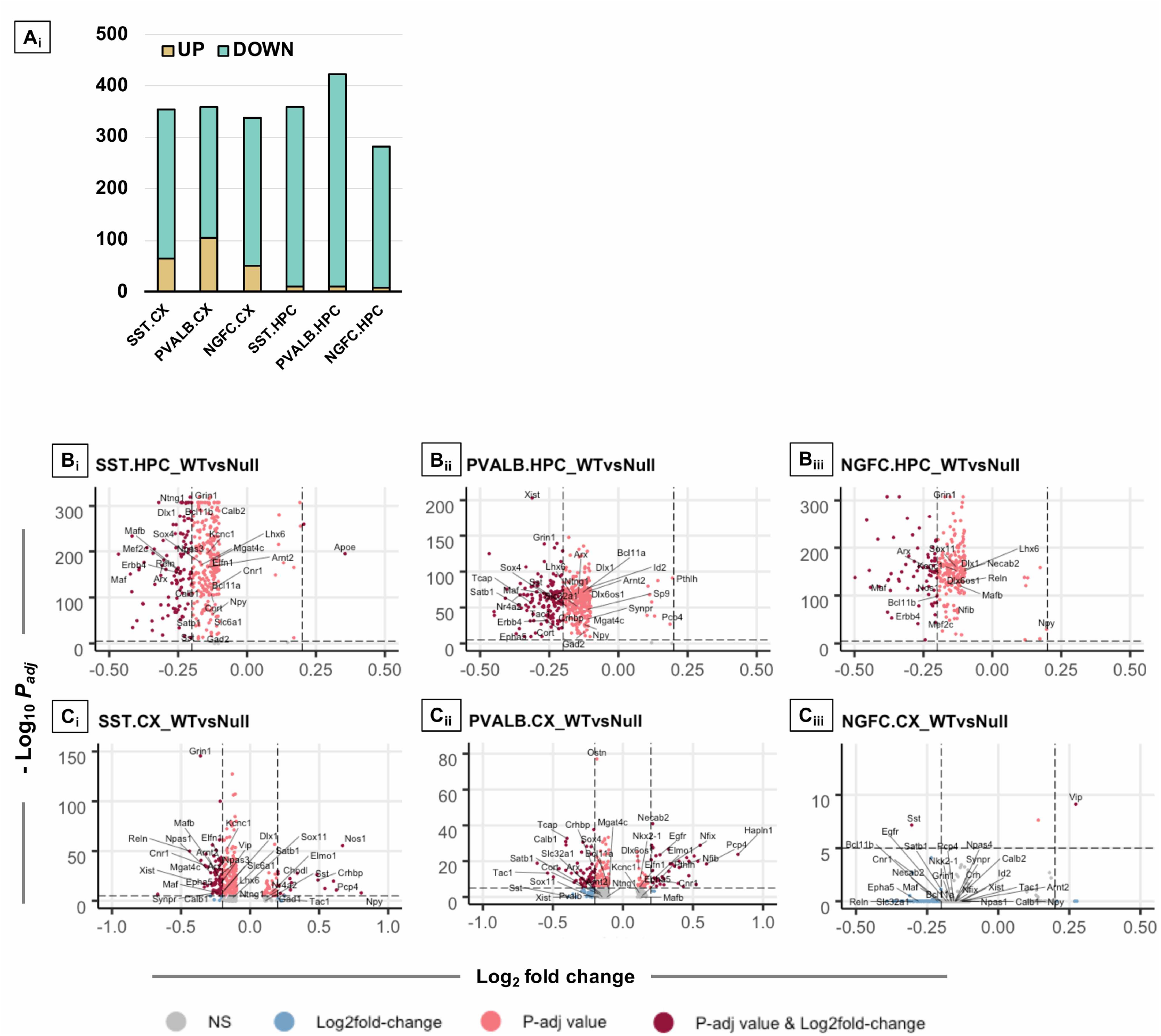
Differential gene expression in the MGE subtypes subsequent to *Grin1*-ablation. **A**_i_, Split-violin plot from both genotypes indicating the expression of *Grin1* in the MGE-derived interneurons, pyramidal neurons and non-neurons. **A** _ii_, Bar plot denoting the number of genes up/downregulated in the cortical and hippocampal MGE clusters. Volcano plot representing the –log10 False Discovery Rate (FDR) versus log2 fold change (FC) between **B,** hippocampal and **C,**cortical MGE cardinal clusters upon *Grin1*-loss, at a fold change ≥0.2 and FDR <10e-6.

**Figure3, Supp.2.**
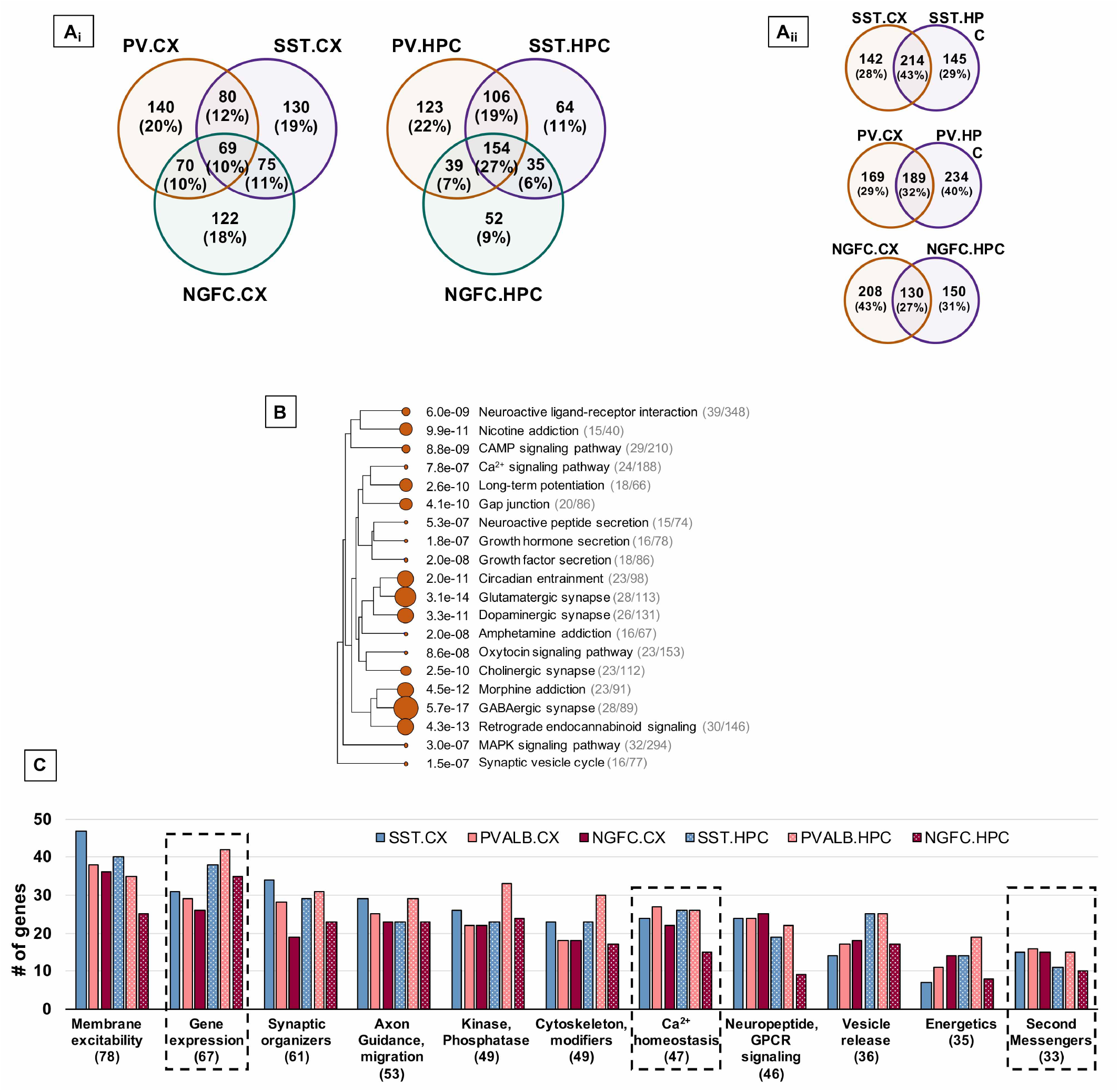
Molecular pathways differentially expressed in MGE subtypes. **A_i_**, Venn-diagrams indicating the percentages of DEGs common within MGE subtypes from cortex or hippocampus. **A_i_**, Venn-diagrams indicating the percentages of DEGs common within MGE subtypes from cortex and hippocampus. **B**, Hier-archical clustering tree summarizing the correlation among significant pathways enriched among the DEGs. Pathways with many shared genes are clustered together. Bigger dots indicate more significant P-values. **C**, Bar plot showing the classification of molecular functions of the DEGs, across the MGE subtypes. Total number of DEGs in the particular molecular class indicated in parentheses.

**Figure4, Supp.1.**
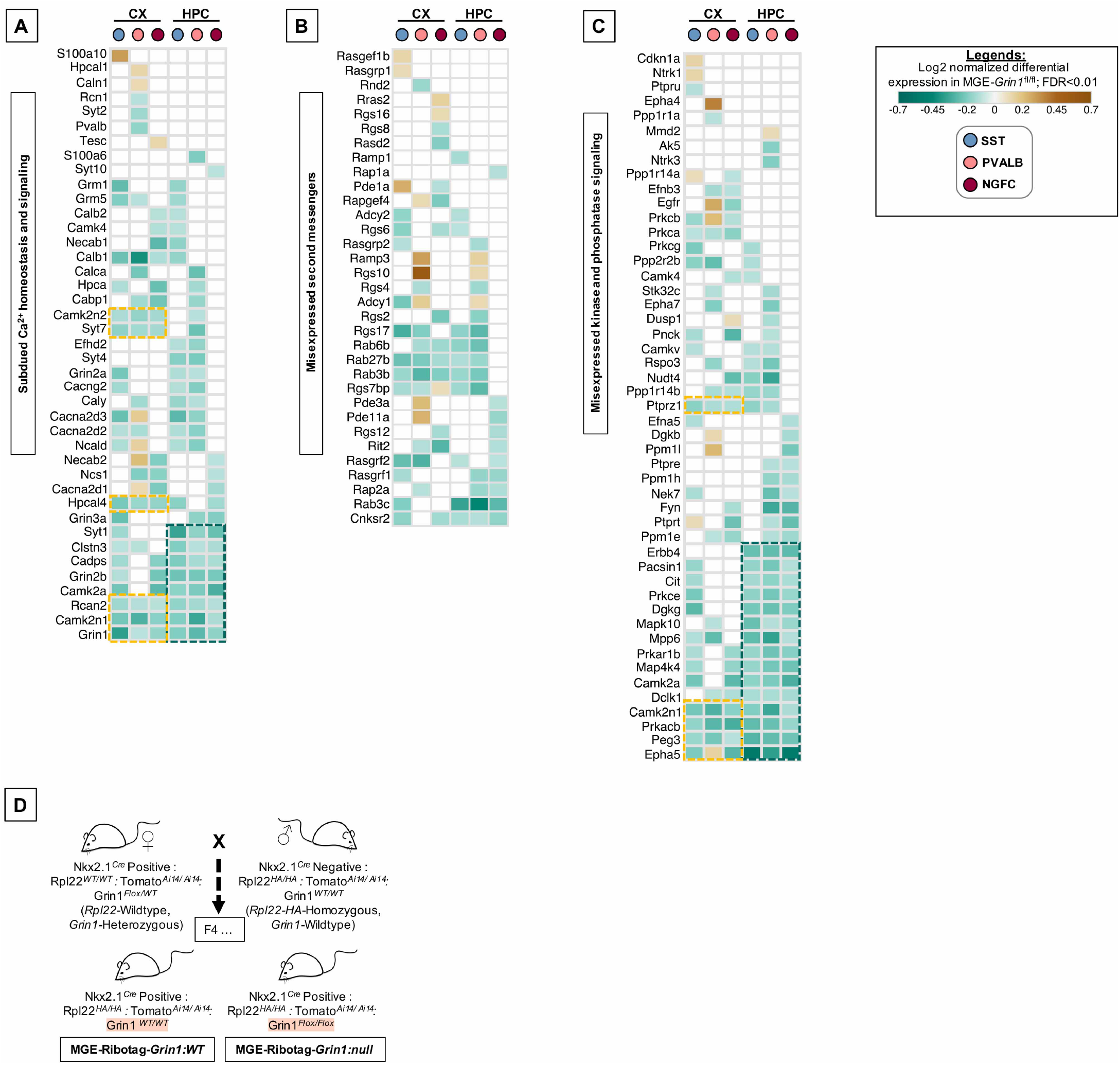
Differential expression of intracellular signaling cascades across subtypes upon *Grin1-* ablation. Heatmap of log2 FC of significant DEGs in cortical and hippocampal MGE cardinal subtypes, showing a subset of **A**, genes regulating intracellular Ca^2+^ homeostasis and Ca^2+^ binding proteins; **B**, notable second messengers; and, **C**, Ca^2 +^ dependent / activated kinases and phosphatases. **D**, Breeding strategy to obtain MGE-derived interneuron-specific expression of Ribotag (Rpl22-HA) transgene in the background of *Grin1^fl/fl^* alleles.

**Figure5, Supp.1.**
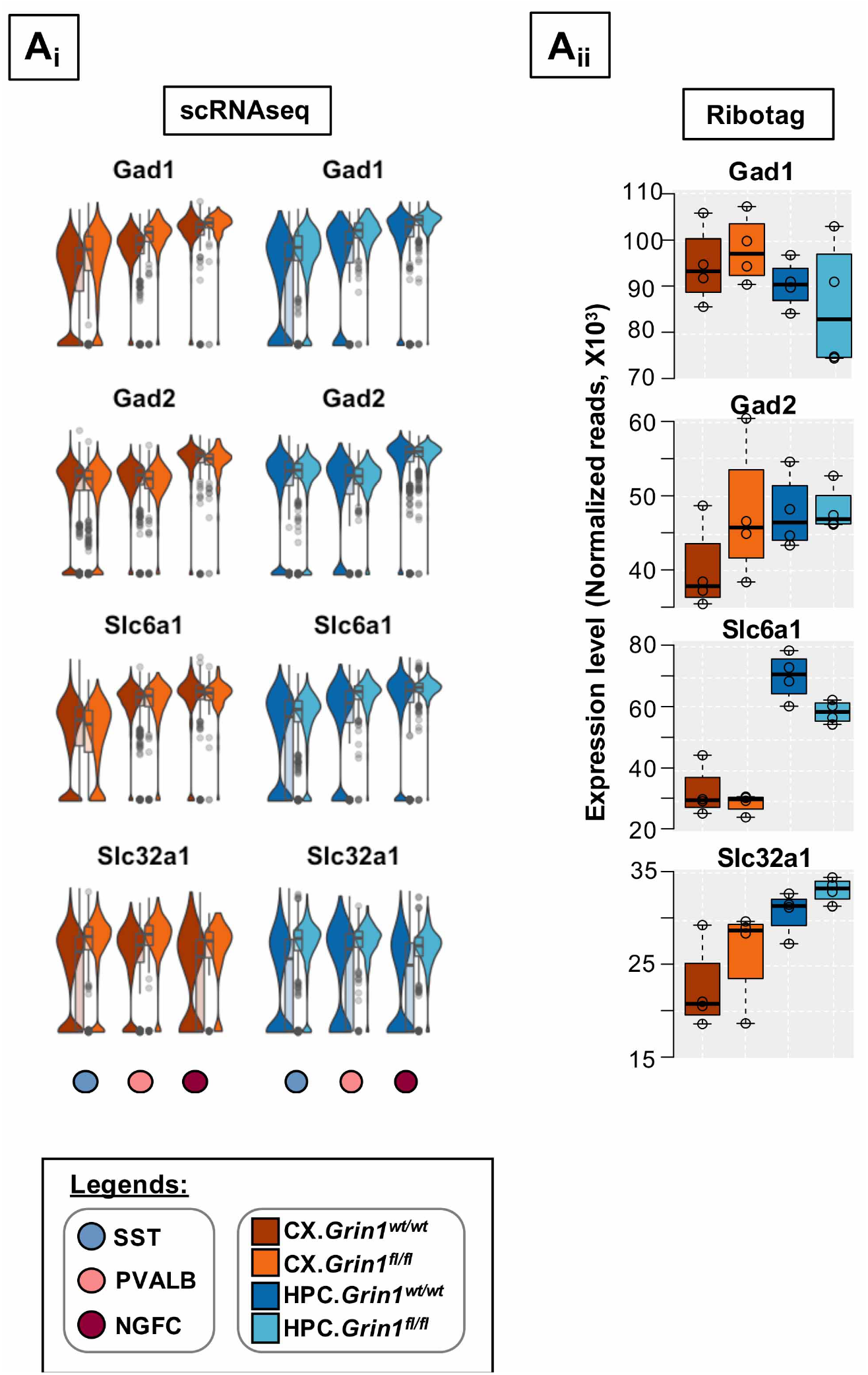
Differential expression of fundamental interneuron marker genes across sub-types upon *Grin1*-ablation. Differential expression of cardinal interneuron marker genes that are broadly expressed across interneurons by **A**_i_, scRNAseq approach, and cross-validated using **A**_ii_, the Ribotag-seq approach.

**Figure6, Supp.1.**
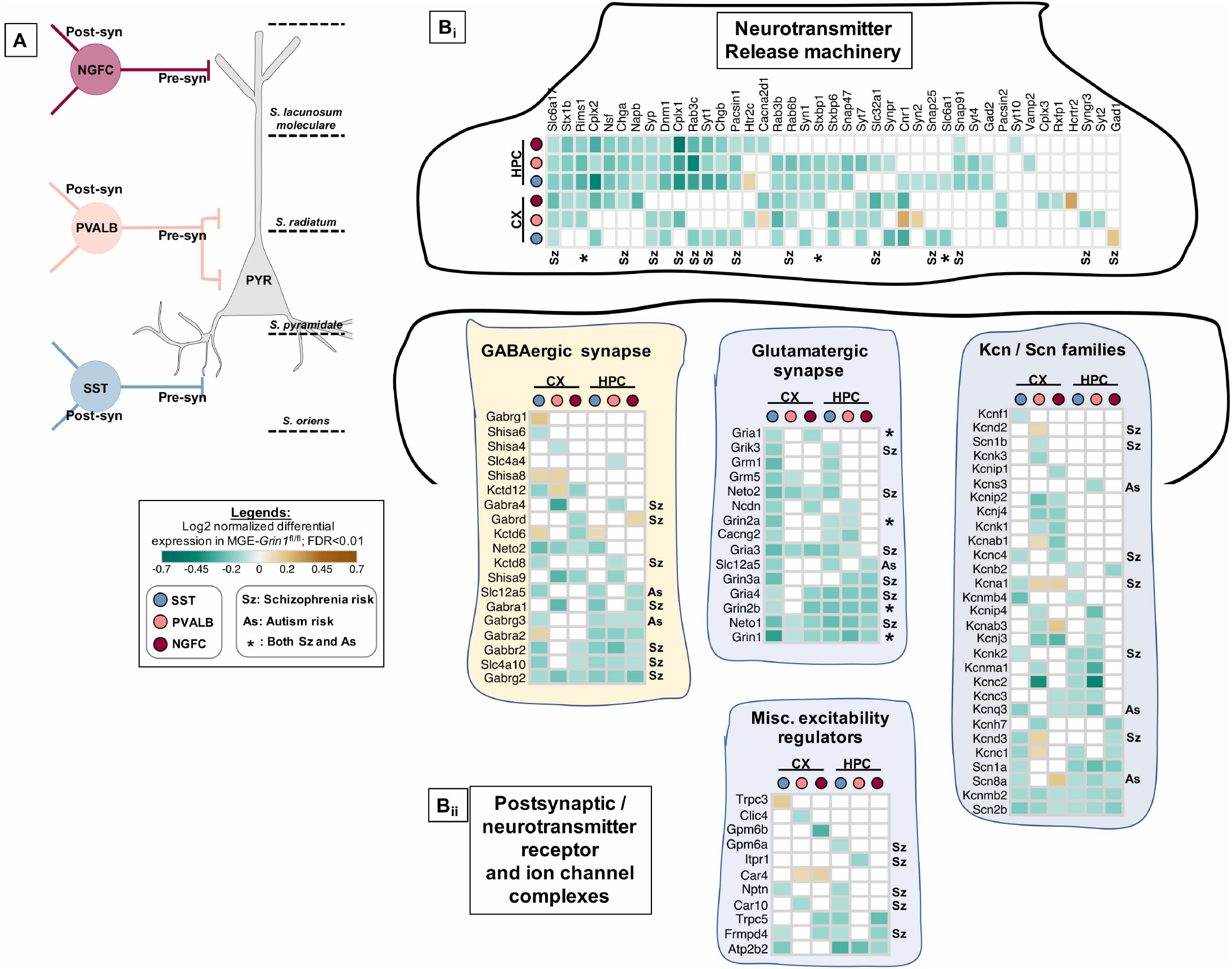
Aberrant NMDAR-signaling result in misexpression of regulators of membrane excitability that are high-risk Sz genes. **A**, Schema representing the field of hippocampal pyramidal cell innervated by the interneurons. Heatmap of log2 FC of significant DEGs in cortical and hippocampal MGE cardinal subtypes, showing a subset of **B_ii_**, Neurotransmitter release machinery. **B_ii_**, Postsynaptic GABA / Glutamate receptor complexes, *Kcn-, Scn-* ion channel complexes and other miscellaneous regulators of excitability.

